# Domestication shaped the chromatin landscape of grain amaranth

**DOI:** 10.1101/2025.04.17.649361

**Authors:** Corbinian Graf, Tom Winkler, Peter J. Maughan, Markus G Stetter

## Abstract

Plant domestication has had profound impacts on the morphology and genetic diversity of crops. Beyond sequence diversity, changes in chromatin structure can play an important role in plant adaptation. However, the interplay between the chromatin landscape and plant domestication remains unclear. We present a high-quality genome assembly and chromatin landscape map of the ancient pseudo-cereal, amaranth. Using ATAC-sequencing of multiple accessions of three grain amaranth species and two wild relatives, we show that the overall amount of accessible chromatin is highly conserved, but about 2.5% of all chromatin switched states, with a higher fraction of the genome repeatedly opening during domestication processes. These differentially accessible chromatin regions, between the crops and their wild ancestor, were species-specific and significantly associated with selective sweeps - reflecting the repeated independent domestication of amaranth. Our findings reveal the dynamic interplay between domestication and the chromatin landscape, highlighting an additional layer of diversity in crops.

## Introduction

Plant domestication has had profound impacts on humans, plants, and even ecosystems [1, 2, 3]. The adaptation of plant populations to the human-made environments required numerous trait changes and imposed selective pressure on plant populations. Some of the traits that changed during domestication have been studied in recent years. These include morphological [4], physiological [5] and intrinsic traits such as metabolite composition [6] and gene expression [4]

While the impact of domestication on numerous traits and genetic diversity has been studied [7], the impact on physical properties of DNA has only recently started to receive attention. Methylation has been shown to vary genome-wide between domesticates and their wild relatives [8, 9] and the change has been associated with several phenotypic changes [10, 11, 12]. In addition, genome size has changed in multiple domesticated species, even in the absence of whole genome duplications [13, 14, 15], and polyploidization has also been associated with species that harbor domesticates [16]. These examples show that the physical properties of the genome interact with selection during plant domestication. However, the extend and repeatability of such changes remains unclear.

The chromatin landscape describes the physical constellation of the DNA molecule packed in the nucleus. DNA can be tightly packed as heterochromatin (closed), or not bound to histone proteins and therefore accessible to other molecules (euchromatin) [17]. The open state of euchromatin allows transcription factors and polymerases to bind to DNA and be transcribed. Through its regulation of DNA accessibility, the chromatin state influences gene expression, local recombination rate, transposable element (TE) activity, and genome structure [18, 19, 20]. While chromatin state changes can be plastic and vary between tissues and even between individual cells [21], heritable chromatin differences are observed on the species, population, and between-individual level [22, 23].

The importance of chromatin for gene expression and other essential functions suggests that chromatin and the chromatin landscape might be involved in adaptation. For instance, the increased capability of *Saccharomyces cerevisiae* and *Dekkera bruxellensis* to ferment glucose under aerobic conditions is caused by changes in the chromatin state of mitochondrial genes, which evolved independently in both species [24]. A few studies have looked at the impact of plant domestication on chromatin accessibility. In soybean a comparative genomic analysis revealed a 3.7% change in chromatin state [23] and approximately 21,000 chromatin loops were differentially formed between wheat (*Triticum aestivum*) and each of its wild relatives *Triticum durum* and *Aegilops tauschii* [25]. However, most studies to date have relied on comparative analysis between just a few accessions of the crop and the wild ancestor, limiting their power to detect chromatin state changes, since the variation can be high even between individuals of the same species [26]. Hence, a population-scale comparison along the domestication gradient should have the power to reveal the interplay between directional selection on morphological traits during domestication and the chromatin landscape.

One model that is well-suited for this purpose are the grain amaranths, *Amaranthus caudatus* L., *Amaranthus cruentus* L. and *Amaranthus hypochondriacus* L.. These nutritious diploid pseudo-cereals from the Americas have been domesticated at least three times from one ancestral species (*A. hybridus*) [27]. The most commonly cultivated species, *A. hypochondriacus* and *A. cruentus*, were domesticated in Central and North America, respectively, while *A. caudatus* was domesticated in South America. The wild *Amaranthus* species, *A. quitensis*, has previously been suggested to have been involved in the domestication of *A. caudatus*. Indeed, strong signatures of gene flow are seen between these sympatric relatives [28]. The replicated domestication and parallel selection for domestication traits make the grain amaranth species complex an ideal system to study the impact of domestication and selection on the chromatin landscape.

We sequenced the pan-chromatin map of 42 samples representing the five *Amaranthus* species that were involved in the repeated domestication of the crop. We first assemble an improved reference genome, its methylome and high-resolution chromatin map through Assay for Transposase-Accessible Chromatin (ATAC) sequencing for the domesticated species *A. hypochondriacus*. We show that a number of transposon families preferentially insert into open chromatin regions, but methylation silences a large proportion of retrotransposons in open chromatin regions. While chromatin accessibility is mostly conserved through the domestication process, a significant number of regions changed their chromatin state during domestication. Consistent with the previously identified independent domestications of the three grain amaranths, most chromatin changes are species specific. Interestingly, several key candidate regions have overlapping selection signals, suggesting an interplay between chromatin and selection during plant domestication.

## Results

### Highly complete A. hypochondriacus reference genome and annotation shed light on genus evolution

To facilitate our genomic analyses, we generated an improved *A. hypochondriacus* reference genome assembly, using a sequencing depth of 30X HiFi (mean length: 13 kb) and 32x ONT (mean length: 48 kb, Supplementary Fig. S1), resulting in a total assembly length of 434,800,201 divided into 348 contigs (N50: 9,806,733 bp, L50: 16). The assembly was further scaffolded using Hi-C data and manually inspected to capture the 16 haploid chromosomes of *A. hypochondriacus*, producing the final 434,863,491 bp assembly consisting of 232 scaffolds (N50: 25,996,252 bp, L50: 7), including the 16 chromosome-level scaffolds, which alone comprise 96.4% of the sequence. The final assembly demonstrated high BUSCO completeness (99.3% complete BUSCOs, including 2.12% duplicated) and an assembly length increase of 31 Mb compared to the previous, primarily short-read based, reference genome [434,863,491 bp compared to 403,994,491 bp; 29, 30]. The new assembly was largely collinear to the previous reference genome, however, chromosome 11 featured a large inversion compared to the previous reference genome (Supplementary Fig. S2). A segment of chromosome 10 that is hypothesized to contain an erroneous assembly of organellar contigs in the previous assembly [30], shows considerable rearrangement in the new assembly. For both, chromosome 10 and 11, capturing of these genomic regions within single contigs indicate correction of the misassembled regions in the new assembly (Supplementary Figs. S3 and S4). The increased length and BUSCO completeness of the assembly in addition to the correction of putative misassembled regions demonstrate the high quality of the assembly.

To annotate genes in the genome assembly, we combined *ab initio* gene prediction (24,583 genes and 27,697 isoforms, 98.8% BUSCO completeness) with a set of high-quality full-length transcripts obtained from isoform sequencing (35,187 transcripts, 94.5% BUSCO completeness). We merged both computational annotation and protein coding transcripts from the full-length transcriptome into the final genome annotation that includes 25,167 annotated protein coding genes with a total of 30,529 annotated isoforms. The new annotation featured the highest annotation completeness for amaranth to date (98.8% BUSCO completeness; Supplementary Table S1). The annotation of 23,155 isoforms was based directly on full-length transcripts from isoform sequencing and, therefore, includes high-confidence annotation of untranslated regions. A total of 219 Mb (50.43%) of the assembly was annotated as transposable elements and tandem repeats (Table S2). Large parts of the genome were annotated as DNA transposons (98 Mb, 22.62%) and retroelements (109 Mb, 25.17%), including a high fraction of LTR elements (97 Mb, 22.33%; supplementary Table S2). Corresponding to previous reports [31], LTRs and LINEs were mostly annotated in gene-sparse regions while DNA transposons were more evenly distributed along the genome (Supplementary Fig. S5).

We calculated the phylogenetic and syntenic relationship among species of the *Amaranthus* genus and with closely related species to study their genome evolution (Extended Data Fig. E1, Extended Data Fig. E2), which reflect previous results by Raiyemo et al. [32]. Peaks in the distribution of synonymous substitutions (K_s_) indicate a single whole-genome duplication in *Amaranthus* (K_s_ ∼0.5) compared to *Beta vulgaris* (K_s_ ∼0.625, Extended Data Fig. E1). We dated the whole-genome duplication to 24.7 Mya (15.5 Mya - 44.9 Mya), closely resembling the previous estimate of 25.56 Mya by Wang et al. [33]. Within the *Amaranthus* genus, genomes were highly co-linear with few large rearrangements (Extended Data Fig. E1).

### The interplay between chromatin state and methylation mediate transposable element silencing and gene expression

We used the improved reference genome to investigate the general structure of the chromatin landscape of amaranth by creating the first genome-wide map of the chromatin landscape of *A. hypochondriacus* through ATAC sequencing (ATACome) and comparing it with other structures of the genome (1A). We sequenced a total of 8 samples from leaf (n=5) and seedling (n=3) tissue from the reference accession PI 558499 (Plainsman), yielding a total of 308,579,996 read pairs. From the 269,667,484 read pairs that uniquely mapped to the *A. hypochondriacus* reference genome (Extended Data Fig. E3), we identified a total of 142,649 accessible chromatin regions (ACRs) using the MACS3 pipeline [34]. Most ACRs were shorter than 1,000 bp (Supplementary Fig. S6), with the largest at 5,174 bp. We removed the two ACRs that were larger than 4,000 bp for downstream analysis, as they might represent false positives and inflate ACR proportions. On average we identified 22,156 ACRs per sample, covering about 11,238,400 bp (2.56% of the reference genome, Supplementary Fig. S7). This fraction is similar to the range identified in other plant species and fits within the expected correlation between genome size and open chromatin [35, 36, Extended Data Fig.E4], suggesting that open chromatin can approximate functional space in the genome [37]. About 87.94% of ACRs physically overlapped between at least two samples (Supplementary Fig. S8). To reduce inherent variation in sequencing chromatin and create a more robust set of ACRs, we fused overlapping ACRs and discarded ACRs that were found only in one sample. This left the completed ATACome with 29,188 unique ACRs covering 23,071,669 bp or about 5.31% of the reference genome, representing the most complete overview of the functional space of amaranth and its genus to date.

Investigating the landscape of these ACRs, revealed a high association of ACRs with genes regardless of tissue (Fig. 1B). In total, 75.99% of ACRs were part of a gene body (intron: 4.32%, exon: 3.67%) or closely associated with it (2kb upstream of transcription start site (TSS): 40.27%, 2 kb downstream of transcription termination site (TTS): 25.76%) (Fig. 1B). The distribution of ACRs among genome features was significantly different from a random distribution, likely reflecting the close association of ACRs with genes and their role in controlling gene expression [38]. Nearly half (49.59%) of the 30,304 ACRs were shared between the two tissues, documenting the within-plant variation in the chromatin landscape. Almost twice as many tissue-specific ACRs were identified in leaf tissue (8,559) than in seedling tissue (4,920), suggesting a higher differentiation of chromatin in leaves, compared to the seedling samples which were comprised of multiple distinct tissues (cotyledon, hypocotyl, roots) (Fig. 1C).

**Figure 1.**
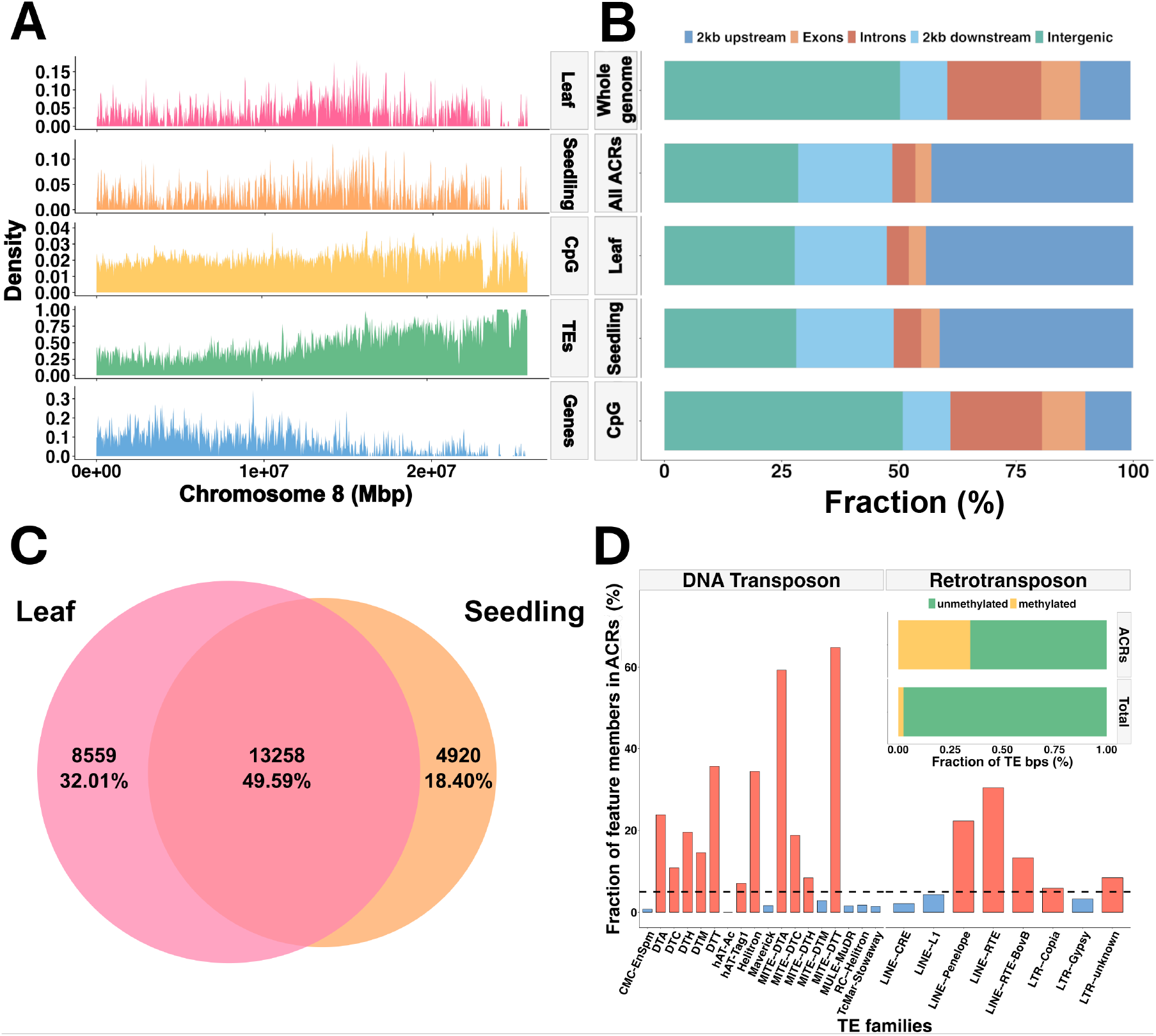
Genome and chromatin landscape of *A. hypochondriacus*. **A**. Overview of the distribution of the transposable content (green), gene content (blue), accessible chromatin content in seedling tissue (orange), and accessible chromatin content in leaf tissue (pink) along chromosome 8. **B**. Distribution of accessible chromatin regions (ACRs) and methylation in CpG context among the genomic regions. **C**. Overlap of peaks called between seedling and leaf tissue samples. **D**. Number of occurrences in ACRs for each of the 26 TE families and whether the family is enriched (blue) or depleted (red) in open chromatin compared to the whole genome. **Inset:** Fraction of methylated (yellow) and unmethylated (green) base pairs for all TEs and TE base pairs within ACRs.

In addition to chromatin, methylation can control functional regions in the genome. As such, we generated a methylation map of *A. hypochondriacus* using methylation data obtained from whole-genome bisulfite sequencing (WGBS) of young leaf tissue from Plainsman using Bismark [39]. Of the 85,240,870 reads, 77,502,531 uniquely mapped to the reference genome. In total, 16.98% of Cs in the genome were methylated. Of these, 8.07% were in a CpG context, 4.63% were in CHG context and 4.28% were in a CHH context. Of the Cs in a CpG context, 75.77% were methylated, while 41.64% of Cs in CHG context and 5.47% of Cs in CHH context were methylated. Methylation in CpG, CHG and CHH contexts were higher in amaranth than observed in *Arabidopsis* but similar to grapevine, which has a similar genome size and TE content as amaranth [10, 40, 41]. About 50.82% of methylation was accumulated in intergenic regions, 9.85% upstream of genes and 10.15% in gene bodies (Fig. 1B). Methylation in ACRs was significantly depleted with only 3.17% of open chromatin being methylated, indicating that the majority of accessible chromatin allows binding. Accordingly, the lowest methylation density was at the TSS, aligning with the highest density of accessible chromatin (21 bp downstream from the TSS) and probably facilitating the expression of genes (Fig. 2A).

**Figure 2.**
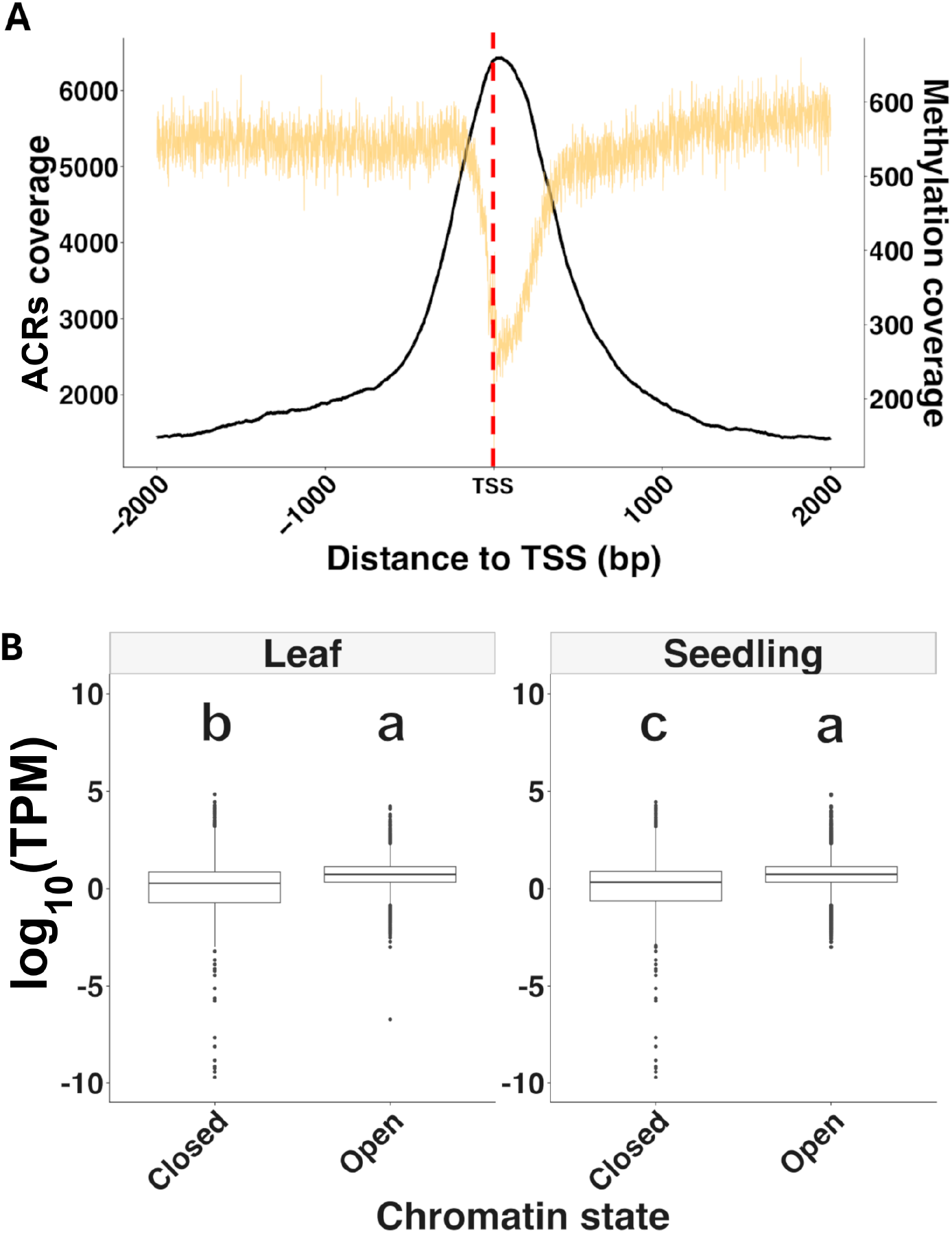
Association between chromatin and gene expression. **A**. Accessible chromatin region (ACR) coverage (black) and methylation coverage (yellow) near the transcription start site (TSS). The red line indicates the TSS. **B**. Mean gene expression across 8 tissues of genes in accessible chromatin regions (ACRs) (open) and equal number of random genes (closed). Letters indicate significant differences between groups based on an ANOVA.

Increased chromatin accessibility upstream of the TSS is often associated with higher expression of the corresponding gene. Indeed, we found that genes associated with tissue-specific ACRs showed significantly higher expression in their corresponding tissues compared to a similar sized set of randomly selected genes which were not associated with ACRs (Supplementary Fig. S9). This was the case for ACR genes in leaves and in seedlings (Fig. 2B). Despite the significantly higher expression of ACR genes, a gene ontology enrichment analysis did not find any significant terms that overlapped between tissues, but 26 enriched functions for seedling ACR genes and 13 enriched functions in leaf tissue ACR genes were identified (Supplementary Fig. S10). In leaf tissue, most enriched functions pertained to metabolic processes, while in seedling tissue ‘signaling’ and ‘response to stimuli’ were the most common.

The interaction of chromatin and transposable elements (TEs) is also important for the functionality of plant genomes. TEs pose a threat to the integrity of the genome, due to their ability to insert into functional regions of the genome, potentially disrupting vital genes or other important regulatory elements [42]. Hence, the suppression of TE transposition by ‘locking’ them into condensed (closed) chromatin, is likely one of the important functions of the chromatin state [43]. While most ACRs (78.83%) overlapped with TEs by at least 1bp, the chromatin landscape was overall depleted for TEs compared to the rest of the genome. Indeed, TEs made up 47.74% of the *A. hypochondriacus* reference genome but only 35.23% of open chromatin (Supplementary Fig. S11). We found 11 DNA-transposon families and five retrotransposon families to be enriched in ACRs (Fig. 1D). This enrichment may result from the preference of these TE families to insert near gene bodies, which positions them within open chromatin and prevents their effective silencing through containment in closed chromatin [44]. Of the 63,782 TEs within open chromatin, 82.31% overlapped partially with ACRs while 15.48% were completely open, another 2.21% of the TEs were large enough to carry entire ACRs with them. Therefore, the majority of ACR associated TEs were not fully accessible and thus potentially inactive. Furthermore, methylation within ACRs was higher in TE sections, where 78.29% of methylated Cs were part of TEs. The number of methylated base pairs of TEs within ACRs was higher than for TEs outside of ACRs (Fig. 1D).

### Repeated enrichment of selective sweeps within differentially accessible chromatin during amaranth domestication

To elucidate the role that adaptation played in the divergence of the chromatin landscape between species, an accurate representation of the diversity within each species is needed. As such, we sequenced a total of 42 samples from leaf (n=22) and seedling (n=20) tissue from 18 accessions across five species, specifically *A. caudatus, A. cruentus, A. hypochondriacus, A. hybridus*, and *A. quitensis*, yielding a total of 1,710,942,952 read pairs (supplementary Table S3).

Genome comparisons are often complicated by the alignment to a single reference that might be unevenly related to the studied individuals. To assess the potential of reference bias, we aligned our *A. hypochondriacus* ATAC-seq data to the *A. cruentus* reference genome [45]. We called a total of 156,378 ACRs on the *A. cruentus* genome, compared to 142,647 when mapping to *A. hypochondriacus*. By joining physically overlapping ACRs we confirmed 30,792 unique ACRs covering 6.5% of the *A. cruentus* reference genome, similar to the 5.29% of the *A. hypochondriacus* reference. The association of ACRs with genes was also seen in *A. cruentus*, where 68.63% of ACRs were found in (exon 5.13%, intron 5.6%) or near genes (2 kb upstream 38.51%, 2 kb downstream 19.39%) (Supplementary Fig. S12). As in *A. hypochondriacus*, nearly half (46.37%) of the unique ACRs were found in both tissues, with 7,579 and 5,746 being unique to leaf and seedling tissue, respectively (Supplementary Fig. S13). Independent t-tests comparing ACRs called from the two reference genomes showed that neither the number of ACRs called (p = 0.4138), the distribution of ACRs along the genome (p = 1), nor the distribution among tissues (p = 0.8822) significantly differed between the different reference genomes. Together, these results suggest no significant impact of reference bias when calling ACRs of different grain amaranth species. Hence, we carried out our multiple species comparisons using the new *A. hypochondriacus* genome.

Across all samples of the five species, 79% of the reads uniquely mapped to the *A. hypochondriacus* reference genome. A principal component analysis (PCA) of single nucleotide polymorphisms called from ATAC-seq data reconstructs the population structure similar to the whole genome sequencing data (Supplementary Fig. S14A). Calling coverage peaks to identify accessible chromatin from this data identified a total of 178,194 ACRs, covering an average of 2.38% to 2.67% of the reference genome (Fig. 3A). A PCA based on ACR presence-absence separated the samples stronger by tissue than by species, contrary to the PCA on genotypic data, but aligning with the important developmental function of chromatin (Supplementary Fig. S14). An ANOVA revealed that there was no significant difference between species in total amount of open chromatin (Fig. 3A) or total number of ACRs (Supplementary Fig. S15). The merging of disparate ACRs from all species resulted in 51,571 distinct ACRs covering a total of 10.2% of the reference genome. ACRs that were shared by all species were the most frequent (31.07%, Fig. 3B), similar to the large overlap between tissues (Fig. 1C). The next most common groups were ACRs unique to each of the three domesticates (5.61% to 10.32%) and unique to their wild ancestor (5.05%). This indicates that most changes within the chromatin landscape that occurred during domestication were species specific (a total of 32.43% of all ACRs), reflecting the separate domestication processes. In addition, nearly 10% of ACRs were shared between two or more domesticates, but not with a wild relative, indicating that a large fraction of the ACRs changed during domestication. To control the observed overlaps of ACRs between species for unequal sample size and structure, we permuted species assignments for every species combination (Fig. 3B). For 29 of the 31 species combinations the ACR count was outside the 95% confidence interval, suggesting that they are not randomly overlapping. Comparing each of the domesticated species (*A. caudatus, A. cruentus*, and *A. hypochondriacus*) to the wild ancestor *A. hybridus*, we identified between 13,856 and 15,742 total ACRs that changed their state (closed-open and open-closed, respectively). These differentially accessible chromatin regions (dACRs) that were potentially altered during the domestication process made up between 2.24% to 2.6% of the reference genome (Fig. 3C). In all three of the domesticated species, more chromatin opened (*A. caudatus*: 1.91%, *A. cruentus*: 1.16%, *A. hypochondriacus*: 1.66%) than closed (*A. caudatus*: 0.69%, *A. cruentus*: 1.08%, *A. hypochondriacus*: 0.69%) during the speciation of the domesticates. In comparison, between the two wild relatives (*A. hybridus* and *A. quitensis*) 0.93% opened and 0.99% closed, differing from the pattern observed during domestication (Fig. 3C). Together, this suggests that during the domestication more chromatin opened than closed, but not between the two wild species.

**Figure 3.**
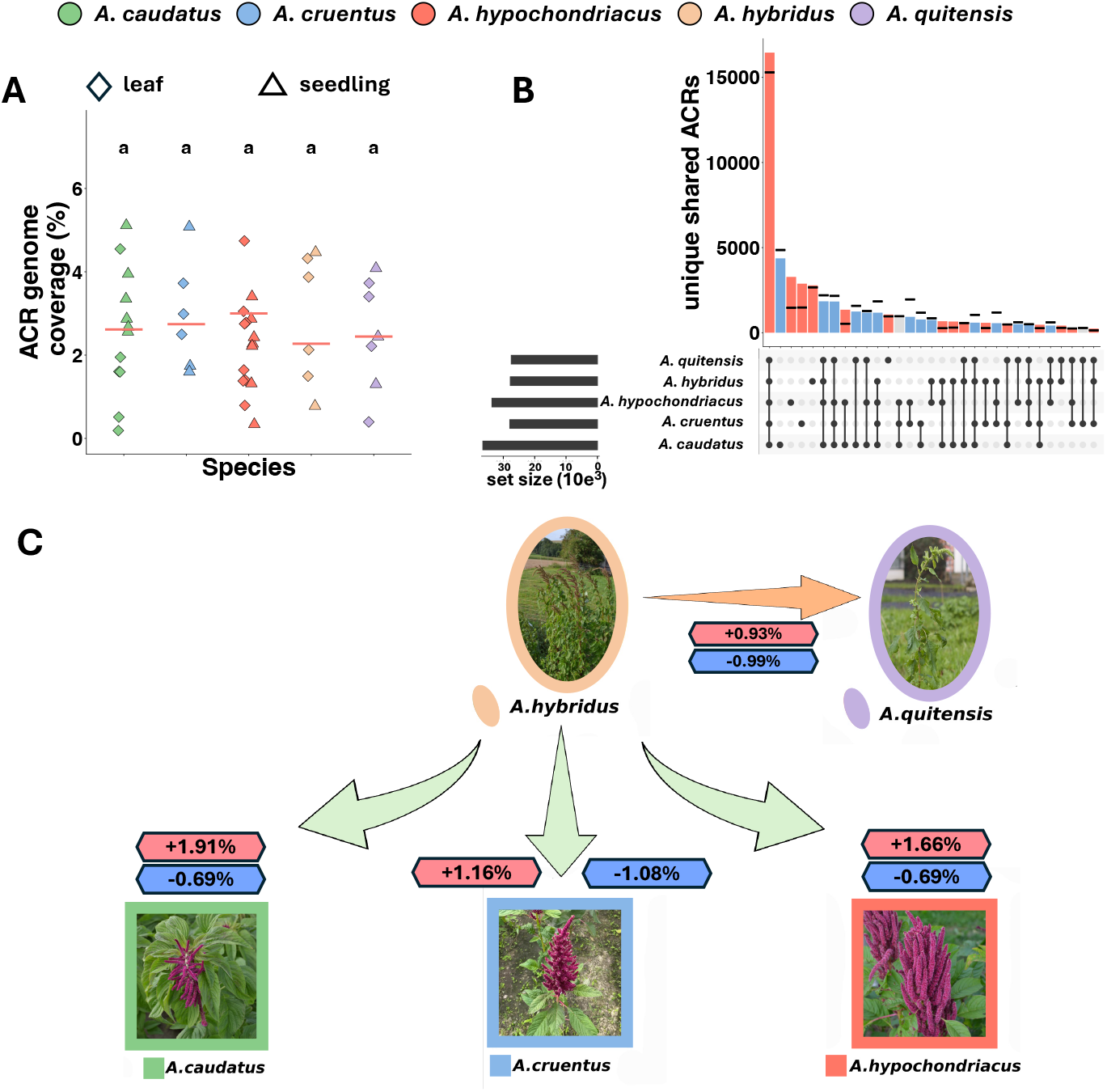
Chromatin changes during amaranth domestication. **A**. Fraction of genome covered by ACRs in each sample. Red lines indicates the mean coverage per species. **B**. Number of unique and shared ACRs across species. Colors indicate whether the groups are enriched (red), depleted (blue) in ACRs or within (gray) the confidence intervals (black lines). **C**. Fraction of genome in differentially accessible chromatin between wild *A. hybridus* and grain amaranth species that opened up (red boxes) or closed (blue boxes) compared to the chromatin landscape of *A. hybridus*

The significant number of dACRs identified suggests that specific chromatin regions might have been under selection during the domestication process (Fig. 4). To test for signals of selection, we analyzed ACR frequency changes between the wild *A. hybridus* and each of the crop species. In total, we found 1,600 dARCs that were fixed closed in the wild ancestor (*A. hybridus*) and fixed opened in the crop species (*A. caudatus*: 645, *A. cruentus*: 726, *A. hypochondriacus*: 229) and 1,293 dACRs that were closed and fixed during domestication (*A. caudatus*: 398, *A. cruentus*: 563, *A. hypochondriacus*: 332)(Fig. 4A). There was no significant overlap between fixed differences (wild-domesticated) across crops species (Supplementary Fig. S16), indicating that selected dACRs were species specific. This agrees with the previous finding based on SNP-based selective sweeps, which also differed among the three domesticated species [27].

**Figure 4.**
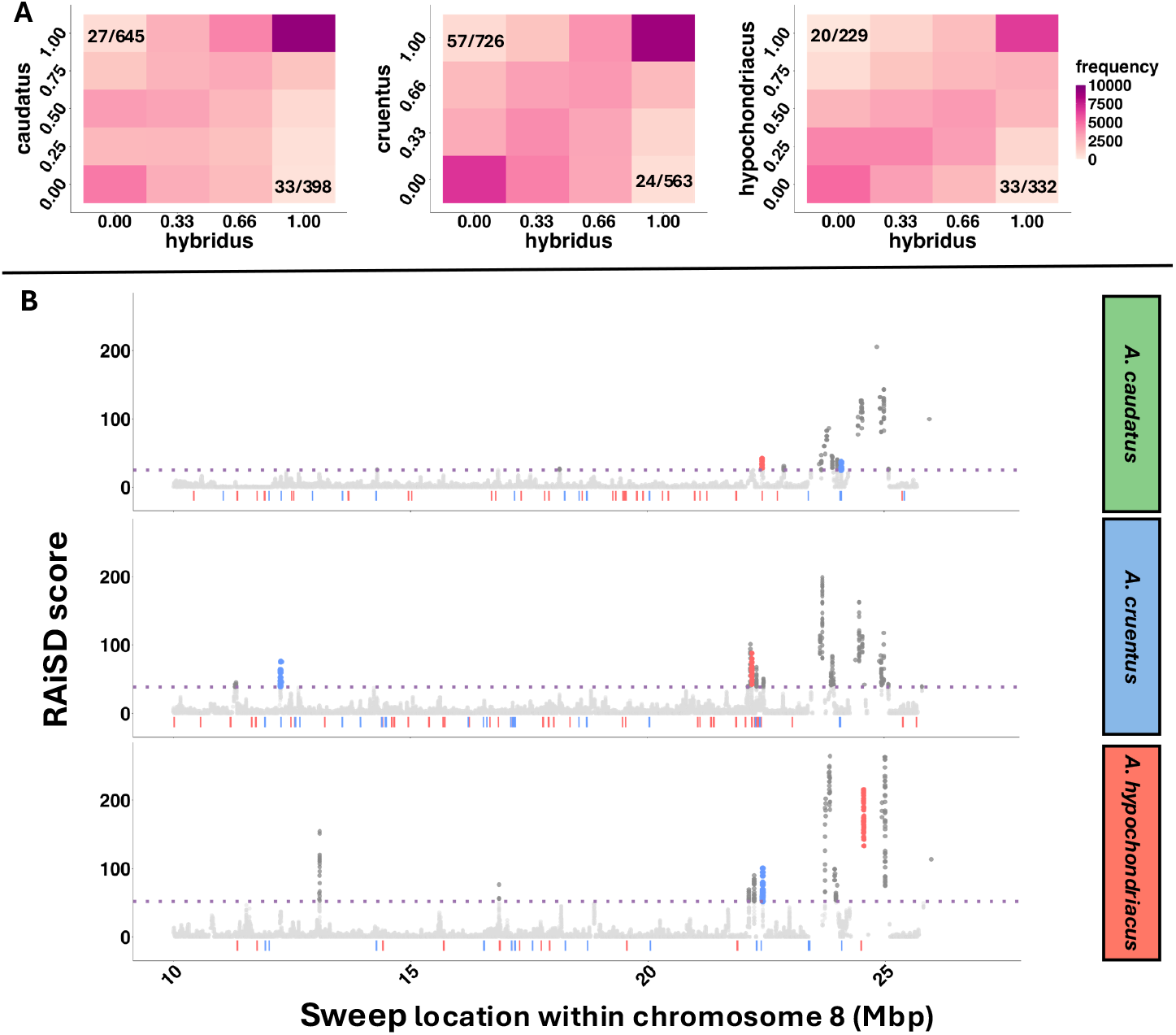
Selection on chromatin during domestication. **A**. Joint frequency of ACRs in domesticates and *A. hybridus*. x and y axis indicate the ACR frequency across accessions in the respective species. **B**. Overlap of fixed dACRs that opened during domestication (red) or closed (blue) and selective sweeps in a subsection of chromosome 8. The purple dotted line indicates the significance threshold for selective sweeps.

To further elucidate the selective role of dARC regions during the domestication, we investigated their association with selective sweeps. We found that between 3.41% to 6.63% of domestication dARCs overlapped with selective sweeps in the respective species (Fig. 4A). Of the dACRs within selective sweeps, 46 were associated with genes (within 2 kb from TSS and TTS). Of these we found six that opened and nine that closed in *A. caudatus*, six opened, six that opened and 11 that closed in *A. cruentus*, and seven that opened and seven that closed in *A. hypochondriacus* (supplementary Table S4). A hyper-geometric test showed a significant enrichment of selective sweeps within dACRs in all three domesticates. While none of the genes that opened during domestication was shared between the species, one of the genes that closed was shared between *A. caudatus* and *A. hypochondriacus*, two were shared between *A. cruentus* and *A. hypochondriacus* (including a disease resistance gene AHq015015) and three that were shared among all three domesticates. One of the genes that closed in all three domesticates was also a disease resistance gene (AHq015012). Two of them belonged to the HAUS augmin-like complex (AHq020586 and AHq020587) coding for the subunit 1 (AUG1), which has an essential developmental function. The potential involvement of the HAUS augmin-like complex in crop domestication and the potential influence of chromatin state on its function should be further investigated through molecular assays.

To investigate the functional link between chromatin accessibility and phenotypic changes, we looked at seedling coloration in *A. caudatus*, a trait with mendelian inheritance [29]. The red pigmentation through betalains is controlled by the expression of the key enzyme CYP76AD2 [30]. We compared the chromatin state between red seedlings and green seedlings and find that upstream of the gene, in the potential promoter region, red seedlings had a stronger accessible chromatin signal than green seedlings (supplementary Figure S17). This connection between dACRs and phenotypic differences illustrates the probable importance of chromatin changes for trait changes. Through dissecting the genetic control of domestication traits in amaranth more links between dACRs and phenotypes can be revealed.

## Discussion

We present a population-scale investigation of chromatin-landscape changes during amaranth domestication. Our study provides the most complete genome assembly, methylome and chromatin map of grain amaranth (*A. hypochondriacus*), enabling functional studies in the orphan-crop and the *Amaranthus* genus as a whole that were previously hindered by the lack of annotation of functional regions and gene models [29, 30, 46]. The quality of our assembly and chromatin map is comparable to that of major crops [23, 35, 36].

Our results highlight how chromatin is integrated into the broader genomic landscape and contributes to the genome ecosystem [47]. We show that approximately two-thirds of ACRs are closely associated with genes across tissues, with about 40% of ACRs associated with the promoter region alone. The high density of ACRs upstream of genes emphasizes the functional significance of promoter regions and illustrates the utility of chromatin profiling in delineating functional genomic elements beyond coding sequences [48, 49]. Nonetheless, ACRs distant to genes might also be part of the functional space, as distal ACRs often contain enhancers, sRNAs and other regulatory regions [49, 50]. Accessible chromatin enables transcription factor binding and gene expression [36], but also increases the proliferation of TEs [44]. While we find a large overlap between TEs and ACRs, most TEs were only partially open (less than 25% of TE in ACRs). Of the TEs that were located in ACRs, a large fraction (32.4%, compared to 4.49% genome-wide) were methylated, which might lead to their silencing even when they were located in an ACR. Whether the over-representation of certain TE families in open chromatin regions results from their preferential insertion into gene-rich areas or is facilitated by the accessibility of the chromatin in those regions remains unclear [47]. Further characterization of the genomic landscape will be critical to understand the interplay between genetic elements and other genomic properties including methylation, chromatin state and recombination rates.

Characterizing the chromatin landscape at the population level provides insights into its role in evolutionary processes [49, 51]. During the domestication of the grain amaranths, the total amount of accessible chromatin remained roughly the same across the five species studied, however ∼2.5% of chromatin changed state in each of the three domestication processes. In all three transitions, a higher fraction of the chromatin opened, indicating a trend towards more open chromatin (Fig. 3C), consistent with findings in soybean [23], maize [52] and rice [9]. Genes associated with ACRs are expected to exhibit higher expression levels compared to non-associated genes [20][(Fig. 2B]. Considering the increased level of open chromatin accessibility, an overall increase in gene expression is expected during domestication. Similar patterns have been seen in other domesticated species, such as maize [53] and cotton [54], in which an increase in overall gene expression has been observed. This effect is particularly pronounced for genes governing traits that were directly under selection [55]. These regulatory modifications are reflected in the histone modification and DNA conformation [23].

The impact of chromatin accessibility changes on phenotypes could drive their selection, either through genomic mutations that lead to altered chromatin or through epigenetic remodeling. Our finding of a significant overlap between dACRs and selective sweeps indicates that chromatin did indeed play a role during domestication. Further research is needed to understand if selection acted on chromatin state with a causal trait change or if the chromatin change was simply a side-effect of mutations but did not alter the traits directly. The group of resistance genes and developmental regulators that we found might be promising candidates for investigating the complex interactions among genomic mutations, chromatin dynamics, trait expression, and selection processes during domestication.

Directional selection during the domestication of plants and animals has long served as model to understand evolutionary changes [56]. Despite the clear impacts of domestication on phenotypes, intrinsic traits, such as the chromatin landscape seem to have responded to selection [57, 58, 59]. This response was either through the direct control of domestication traits by the chromatin state or through a pleiotropic relationship between phenotypes and chromatin state. Assessing the diversity of chromatin and their functional importance could unlock an additional set of variation for crop improvement.

## Acknowledgments

We thank Roswitha Lentz and Christoph Göttlinger for their help with ATAC-library preparation. We acknowledge funding by the Deutsche Forschungsgemeinschaft (DFG, German Research Foundation) under Germany’s Excellence Strategy – EXC-2048/1 – Project ID 390686111 and grant STE 2654/5 to MGS by the DFG.

## Data availability

All generated sequencing data is available through the European Nucleotide Archive (ENA) under the project number PRJEB88670. The newly assembled *A. hypochondriacus* genome and annotation files are accessible at https://quinoadb.org/download/Amaranthus/A.hypochondriacus_Plainsman/. Scripts used in the analyses are available through GitHub (https://github.com/CorbinianGraf/chromatin_landscape_amaranth_2025).

## Online Methods

### Genome assembly and annotation

High molecular weight (HMW) DNA was extracted from fresh leaf tissue of *A. hypochondriacus* (PI 558499; cv. “Plainsman”) using a CTAB-Genomic-tip protocol as described by Vaillancourt and Buell [60]. The HMW DNA was quantified with the Nanodrop^™^ One/OneC Microvolume UV-Vis Spectrophotometer (Waltham, MA), and screened for quality control parameters including DNA concentration (> 800 ng/mL) and contamination (260/280 and 260/230 ≅ 2.0). For PacBio HiFi sequencing, HMW DNA was sheared to 17 kb on a Diagenode Megaruptor (Denville, NJ) and then made into SMRTbell adapted libraries using a SMRTbell Express Template Prep Kit 2.0 (Pacific BioSciences, Menlo Park, CA) followed by size selection using a Sage Science BluePippin (Beverly, MA) for fragments greater than 10 kb. Sequencing was done at the Brigham Young University DNA Sequencing Center (Provo, UT, USA) using Sequel II Sequencing Kit 2.0 with Sequencing Primer v5 and Sequel Binding kit 2.2 for 30 h with adaptive loading using PacBio SMRT Link recommendations. For Oxford Nanopore Technology (ONT) sequencing, HMW DNA was sequenced using the default protocol for ultra-long DNA sequencing kit (SQK-ULK114; Oxford Nanopore Technologies, Oxford, UK) following the standard manufacturer’s protocol with a starting volume of 750 µL (250 ng/µL) on a R10.4.1 flow cell (FLO-MIN114) for 24 hours. Flow cells were washed and reloaded three times. Dorado v0.5.0+0d932c0 with the model ‘dna_r10.4.1_e8.2_400bps_sup@v4.2.0’ was used for base calling of POD5 files. Chopper v0.5.0 [61] was used for trimming (< 30 kb) and quality control (qv < 8.0). In total, approximately 30X coverage of both HiFi (N50 = 14 kb) and ONT reads (N50 = 49 kb) were generated and used for the primary contig assembly.

A primary contig assembly was generated using hifiasm v0.19.8 [62] with default parameters for an inbred species (-l0). Scaffolding of the primary assembly into pseudo-molecules was accomplished using publicly available Hi-C (high-throughput chromosome conformation capture) data (SRA accession: SRR5345531). Hi-C reads were aligned to the primary contig assembly using the Burrows-Wheeler Aligner [63] following the Arima-HiC mapping pipeline (A160156_v03, Arima Genomics, Carlsbad, California, USA). Only uniquely aligned paired-end reads were retained for downstream analyses. Contigs were then clustered, ordered, and oriented into scaffolds using YaHS [64] followed by manual inspection and correction using JuiceBox [65]. The corrected scaffolded assembly was evaluated and corrected with Inspector [66] to produce the final assembly. We named the assembled pseudochromosomes according to scaffold names of *A. hypochondriacus* reference genome v2 [29, 30] and ensured collinearity with the previous assembly version. To prepare the genome for genome annotation, we generated a repetitive element database using RepeatModeler [67] and softmasked the assembly using Repeatmasker [68].

To annotate the *A. hypochondriacus* assembly v3, we combined an *ab initio* gene prediction generated using BRAKER3 v3.0.8 [69] guided by protein and RNA-seq evidence with full-length PacBio Iso-Seq transcripts. As protein database, we used the protein sequences of 117 embryophyta species from ODB10 [70] in addition to annotated proteins of *A. cruentus* [45]. As RNA-seq evidence, we used a public dataset of 31.7 Gb of 90 bp paired-end HiSeq Illumina sequencing data from eight different tissues [BioProject Accession PRJNA263128; 71]. To aid the alignment of RNA-seq reads, we generated a preliminary BRAKER3 gene prediction guided only by protein evidence. We mapped the RNA-seq reads to the assembly with STAR v2.7.8a [72] using the preliminary gene prediction as splice-junction database. We then generated a second BRAKER3 gene prediction guided by both protein and RNA-seq evidence.

For the assembly of full-length Iso-seq transcripts, we used two datasets, from multiple tissues of accession PI 558499: We used a total of 5,3 million circular consensus sequencing (CCS) reads (mean length: 1,441 bp) of public data from seven tissues [root, cotyledon, flower, leaf, pollen, developing seed and mature seed, BioProject ID PRJEB65083; 30]. We further generated 4.4 million CCS reads (mean length: 1,711 bp) from root, leaf, stem, inflorescence and whole seedling tissues of the *A. hypochondriacus* reference accession Plainsman (PI 558499). RNA samples were extracted from each tissue type independently using the Zymo Research (Tustin, CA, USA) Direct-zol RNA MiniPrep Plus kit. The quantity and quality of extracted RNA were evaluated for quality using a Bioanalyzer 2100 (Agilent Technologies, Santa Clara, California, USA). After quality check, RNA from each of the different tissues was pooled in equal molar ratios to synthesize full length complementary DNA (cDNA) using a NEBNext® single cell/low input cDNA synthesis and amplification kit (E6421L), which uses a template switching method to generate full-length cDNAs (New England BioLabs, Ipswich, MA, USA). IsoSeq libraries were prepared from the cDNA of each species according to standard protocols using the SMRTbell v3.0 library prep kit (Menlo Park, CA, USA) and sequenced on a single SMRT cell 8M for each species using a PacBio Sequel II at the DNA sequencing center at Brigham Young University (Provo, Utah, USA).

For both datasets, we processed Iso-Seq CCS reads using Isoseq3 (https://github.com/PacificBiosciences/IsoSeq) and clustered full-length non-chimeric (FLNC) reads into transcripts using Isoseq3 *cluster*. To further deduplicate both datasets, we mapped clustered transcripts to the genome assembly using minimap2 v2.26 [73] with the *splice:hq* preset and collapsed them into sets of unique transcripts using cDNAcupcake 28.0.0 (https://github.com/Magdoll/cDNA_Cupcake) with minimum cutoffs for coverage 0.95 and identity 0.9 resulting in 53,150 and 57,731 transcripts from the public and newly generated datasets, respectively. We used the *chain_samples*.*py* script of cDNAcupcake to combine both datasets, allowing for a maximum 3’ end difference of 300 bp and a fuzzy splice junction distance of 5 bp. We used SQANTI3 v5.2 [74] to correct the combined transcript sequences based on the sequence of the genome assembly. To filter potential artifacts from the combined transcript set, we required transcripts not found in the BRAKER3 prediction to be supported by at least 5 CCS reads and at least 5% of CCS reads per locus resulting in 35,187 transcripts (median length: 1,802 bp, 94.5% BUSCO completeness). To combine full-length transcript sequencing data and BRAKER3 prediction, we predicted open reading frames in the filtered transcript set using GeneMarkS-T [75] and merged both datasets using TSEBRA v1.1.2.4 [76]. We assessed completeness of the Iso-Seq transcripts and the final annotation using BUSCO v5.2.2 [77] in protein mode against the Embryophyta reference set of OrthoDB v10 [70]. For functional annotation of annotated genes, we submitted the annotated protein sequences to Mercator4 v6.0 [78] and eggNOG-mapper v.2.1.12 [79]. To annotate repetitive elements in the genome assembly, we ran the annotation pipeline EDTA v2.2.1 [80] with the *sensitive* parameter and included annotated CDS for purging of gene sequences from the TE library with the *cds* parameter.

### Phylogeny, whole genome duplication and synteny analysis

An orthogroup analysis was constructed with Orthofinder2 v.2.5.4 [81] from longest protein-coding gene models for six *Amaranthus* species with fully assembled genomes (*A. tricolor* [33], *A. tuberculatus* [82], *A. palmeri* [82], *A. retroflexus* [82], *cruentus* [45] and *A. hypochondriacus*) and two outgroup species within the family Amaranthaceae, *Beta vulgaris* L. [83] and *Chenopodium quinoa* Willd. [84]. Genomes for *A. tuberculatus, A. palmeri* and *A. retroflexus* were aquired from the WeedPedia Database [85]. Orthofinder2 assigned 95.1% of all genes to an orthogroup, with a G50 and O50 of nine and 6,919, respectively. A rooted species tree phylogeny was produced using the multiple sequence alignment approach of OrthoFinder2, elicited with the “-M msa” option to produce bootstrap values. WGD v.2.0.38 [86] was employed to identify whole-genome duplications (WGDs) and speciation events by leveraging synteny inference and heuristic peak detection with 95% confidence intervals derived from anchor gene pairs. Synonymous substitution rates (Ks) were calculated between paralogous gene pairs to infer WGDs and between orthologous gene pairs to infer speciation events within and among species. Divergence times among species and WGD events were inferred using a synonymous substitution rate for Amaranthaceae of 9.6E-9 as reported by Wang et al. [33]. Wang et al. reported the divergence between A. tricolor and vulgaris at a Ks peak of approximately 0.63 (divergence between Amaranthoideae and Chenopodioideae subfamilies) and that A. tricolor diverged from B. vulgaris approximately 32.81 Mya. That translates into an estimated substitution rate of 9.6E-9. Divergence time (Mya) = [Ks/(2^*^Substitution rate)]^*^10-6. Syntenic relationships across the *Amaranthus* species were visualized in SynVisio (https://synvisio.github.io/#/) from orthologous genes in collinear blocks, identified with blastp and McScanX [87] using the parameters “-e 1e-50 -k 25 -s 50”.

### Methylation sequencing

Whole genome bisulfite sequencing (WGBS) was generated from young leaf tissue from a single plant from the *A. hypochondriacus* accession Plainsman (PI 558499). Tissue was collected, freeze-dried, and DNA extracted using a modified mini-salts extraction protocol [88]. Quality control parameters for concentration (> 300 ng/mL) and contamination (260/280 and 260/23 ≅ 2.0) were followed before sequencing. The DNA samples were sent to Novogene Corporation, Inc. (San Diego, CA) for WGBS. In brief, the genomic DNA (spiked with lambda DNA) was fragmented to 200-400 bp and then subjected to bisulfite to generate single strand DNA using a EZ DNA Methylation Gold Kit (Zymo Research; Irvine, CA). During the bisulfite treatment, unmethylated cytosine are converted into uracil, while methylated cytosine remained unchanged. Methylation sequencing adapters were ligated, followed by double strand DNA synthesis using the Accel-NGS Methyl-Seq DNA Library kit (Swift Biosciences, Ann Arbor, MI). The quality of the library was verified with Qubit and real-time PCR and the size distribution was verified on a bioanalyzer. Libraries were pooled and sequenced on Illumina NovaSeq X Plus (PE150) instrument to produce a minimum of 30X coverage according to genome size.

We aligned the methylation data to the new reference genome by running Bismark [39] with following parameters “–bowtie2 -n 0 -l 20 methylation/”. The reference genome methylation index was prepared using the bismark_genome_preparation function. The final summary of aligned ACRs was performed using bismarks methylation extractor function. Lastly, we calculated methylation density 2,000 bp up- and downstream from the TSS using an in house script.

### Chromatin accessibility

In order to assess chromatin accessibility changes during domestication, we performed Assay for Transposase-Accessible Chromatin (ATAC) sequencing of a total of 42 samples. These consisted of 3 accession of *A. cruentus* L., 4 accessions of *A. caudatus* L., 4 accessions of *A. hypochondriacus* L., 3 accession of the wild ancestor *A. hybridus* L., and 4 accessions of the wild (potentially feral) *A. quitensis* Kunth. (supplementary Table S3). We sampled leaf tissue from plants grown for 32 days under short-day conditions (8h light, 16h dark) at 22 °C and seedlings tissue from plants grown on filter paper in petri dishes in the dark for 7 days for nuclei extraction.

We extracted nuclei from approximately 50 mg leaf tissue from the fourth fully developed leaf and 100 mg of whole seedlings, according to Wang et al. [89] with minor alterations. In brief, tissue samples were placed in 0.5 ml of chilled lysis buffer (15 mM Tris-HCl pH 7.5, 20 mM NaCl, 80 mM KCl, 5 mM Dithiothreitol (DTT), 0.5 mM Spermine, 1x Cocktail (ThermoFischer: 78429), 0.2% Triton X-100) and finely chopped with a razor blade to release nuclei. The nuclei suspensions were stained with 2 *µ*l of 1 mg/ml 4,6-Diamidino-2phenylindole (DAPI). After confirming the presence of intact nuclei under a fluorescence microscope at 20x magnification, we selected 50,000 intact nuclei based on their size and intensity of DAPI signal under 488 nm excitation using a FACSVantage SE (Beckon Dickinson) and collected them in 0.5 ml lysis buffer. We prepared the 50,000 nuclei for tagmentation by centrifuging them for 4min at 1,000g and 4ºC and discarding the supernatant. Afterwards, we washed the pellet in 1 ml of wash buffer (10 mM Tris-HCl pH 8.0, 5 mM MgCl2, 1x Cocktail) and centrifuged for 4 min at 1,000g and 4°C and discarded the supernatant. We then mixed the nuclei with 25 µl of Tagment DNA Buffer and 2.5 µl TN5 (Illumina: 20034917) and incubated them at 37 °C for 30 minutes. The tagmented samples were purified using the QIAGEN MinElute kit (QIAGEN: 28004). The prepared libraries were sequenced for a minimum of 10 million 100 bp paired-end reads per library on a NovaSeq 6,000 platform by the Cologne Center for Genomics (CCG).

### ATAC-seq data processing

We aligned the sequencing reads to our new *A. hypochondriacus* reference genome using default parameters of bwa-mem2 (2.2.1) [90]. Duplicate reads were identified using picard (2.27.5) and removed with the REMOVE_DUPLICATES=TRUE setting [91]. To exclude overrepresented regions in the genome, we employed a read depth cutoff of 250. Accessible Chromatin Regions (ACRs) were called using the callpeak function of macs3 (v3.0.0) [34] in BAMPE mode with a reference genome size of 439 Mb and a false discovery rate of 0.01. We excluded peaks that only occurred in one library per species and that were longer than 4,000 bp from further analyses as potential false positive calls. To normalize the peak calling for mapability, we also called peaks for whole genome sequencing (WGS) data of each of the 18 accessions [27] and removed ACR peaks from our ATAC-seq data that were also called in the whole genome sequencing data.

### ATACome for PI 558499

We assembled a high confidence ATACome for *A. hypochondriacus* from 8 samples of the reference accession PI 558499 consisting of 5 leaf and 3 seedling tissue samples. In addition to the filters above, we only considered ACRs, which could be called in at least two of the eight samples, to reduce false positives.

We overlapped the ACRs genome-wide with other genomic features, i.e., gene density, transposable elements (TEs) and methylation signatures using the genomicDensity function from the circlize package [92]. Genomic feature annotation of ACRs was performed using the assignChromosomeRegion function of the ChIPpeakAnno package in R [93]. Analysis of the distribution of genomic features for the whole reference genome using assignChromosomeRegion were performed using non-overlapping 10 bp windows due to vector size constraints in R studio. Boundaries were set to 2,000 bp for the upstream (upstream=2,000, downstream=0) of TSS and downstream (upstream=0, downstream=2,000 of TES) genomic features of assignChromosomeRegions. We discarded ACRs that were called in less than two samples of the same tissue to study tissue specific ACRs. We investigated overlaps between tissues with the findOverlapsOfPeaks function of the ChIPpeakAnno package [93] and identified tissue specific ACRs. ACR density 2,000 bp up and downstream of the TSS was calculated using a custom script https://github.com/CorbinianGraf/chromatin_landscape_amaranth_2025. To investigate a potential enrichment of gene functions in open genes, we performed a GO-enrichment analysis, employing the goseq package in R [94]. Peak-length distribution for both tissues was calculated using a custom script https://github.com/CorbinianGraf/chromatin_landscape_amaranth_2025. We compared the expression of open (associated with ACRs unique to either leaf or seedling tissue) and closed genes We compared gene expression between genes associated with open (ACR within 2000 bp of TSS) and closed chromatin within and between tissues, by quantifying gene expression in eight different tissues from the previously described dataset by Clouse et al. [71] using kallisto v0.48.0 [95], calculating the mean expression for each gene across the eight tissues and performing an ANOVA to test for significant differences and corrected for multiple testing using Tukey’s HSD. The expression values were transformed using log10(TPM+0.001). Only genes that were associated with ACRs unique to leaf or seedling tissue were considered. To prevent biases by different sized datasets the smallest set of genes with expression data was identified among the two tissues and chromatin states (open seedling with 10,473 ACR associated genes) and the same amount of genes was randomly sampled from the other three sets. The same conditions were used to compare expression data between chromatin state in each of the eight tissues from Clouse et al. (2016), with the smallest set of genes with expression data being We then determined all genes with ACRs within the gene body or their promoter region (2,000 bp upstream), using the annotatePeakInBatch function of ChIPpeakAnno (output=overlapping, maxgap=2,000) [93]. The gene expression of ACR-associated genes (open genes) was then compared to the expression of an equal number of random genes without an associated ACR (closed genes) using an ANOVA. To study the association between TEs and accessible chromatin, we overlapped ACRs with our TE annotation using the annotatePeakInBatch function from the ChIPpeakAnno package [93]. Enrichment and depletion of TE families in accessible chromatin was determined by performing hypergeometric tests in R using the phyper function from the stats package [96]. Lastly, we calculated the fraction of methylation within ACRs, TEs and TEs within ACRs. Enrichment of methylated base pairs within each of these three groups was tested by performing a hypergeometric test.

### Testing for reference bias

To explore whether aligning ATAC-seq data of multiple species to the *A. hypochondriacus* reference genome causes a bias in peak calling towards data from *A. hypochondriacus* accessions, we also aligned all ATAC-seq data to the *A. cruentus* reference genome [45]. These alignments were subject to the same data processing parameters as for *A. hypochondriacus* aligned data, except for adjusting the genome size during peak calling to the 370.9 Mb of the *A. cruentus* assembly [45]. A chi-square test was performed to test whether the ratio of called peaks between the species is the same between the two alignments. Additionally, chi-square tests were performed to identify if the peak distribution between the two tissues or between genomic regions significantly differed between the two alignments.

### Species comparison of open chromatin

To compare ACRs between species, we used only ACRs that occurred in at least two samples of a species regardless of tissue. Overlapping ACRs from different samples within a species were joined with the reduce function of the GenomicRanges package [97], to create five deduplicated lists of all ACRs within each species. To uniquely index ACRs across species, we combined the species-specific lists into one with unique ACRs for the *Amaranthus* family, by joining ACRs that overlapped between species with the reduce function of the GenomicRanges package. These IDs were then assigned to corresponding ACRs within each species-set. We determined ACR changes in pairwise comparisons between species, based on IDs. To test for statistical significance, we randomized the species identity of each ACR for each sample using the sample function in R and inferred the distribution of ACRs. We calculated confidence-intervals from 100 permutations, using the t.test function of the stats package [98] in R. Differential accessible chromatin regions (dACRs) for each of the three domesticated species compared to the wild *A. hybridus* were identified through a custom script https://github.com/CorbinianGraf/chromatin_landscape_amaranth_2025 and enrichment of gene functions in dACRs was tested using the goseq package.

### Overlap between selection signals and accessible chromatin

We called selective sweeps for each of the five species based on whole genome sequencing data of 28 *A. caudatus*, 21 *A. cruentus*, 18 *A. hypochondriacus*, 9 *A. hybridus* and 12 *A. quitensis* accessions, respectively, taken from Gonçalves-Dias and Stetter [99]. We used RAiSD [100] to identify potential selective sweeps with a cut-off at 0.01% of the highest RAiSD scores. The overlap between selective sweeps and dACRs was determined using bedtools intersect [101]. The enrichment analysis on the selective sweep associated dACRs was performed using goseq. An enrichment of selective sweeps in dACRs was confirmed by testing the overlap of bp in selective sweep regions within ACRs compared to the whole genome, using a hypergeometric test [96].

### Chromatin accessibility near key pigmentation gene

We calculated mean depth for 10 kb up and downstream of CYP76AD2 (chromosome 16: 5238629-5258629 bp) from our ATAC-seq seedling samples of three *A. caudatus* Accession PI 490518, PI 608019, PI 642741 with red seedling color and PI 490612 with green seedling color, respectively. The read count was normalized by dividing the depth through the average depth over the 10 kb. The normalized depth was then overlapped with the 41,812 ACRs found within *A. caudatus*. We plotted the mean normalized depth along the region to examine regions that are differentially open.

## Extended figures

**Figure E1.**
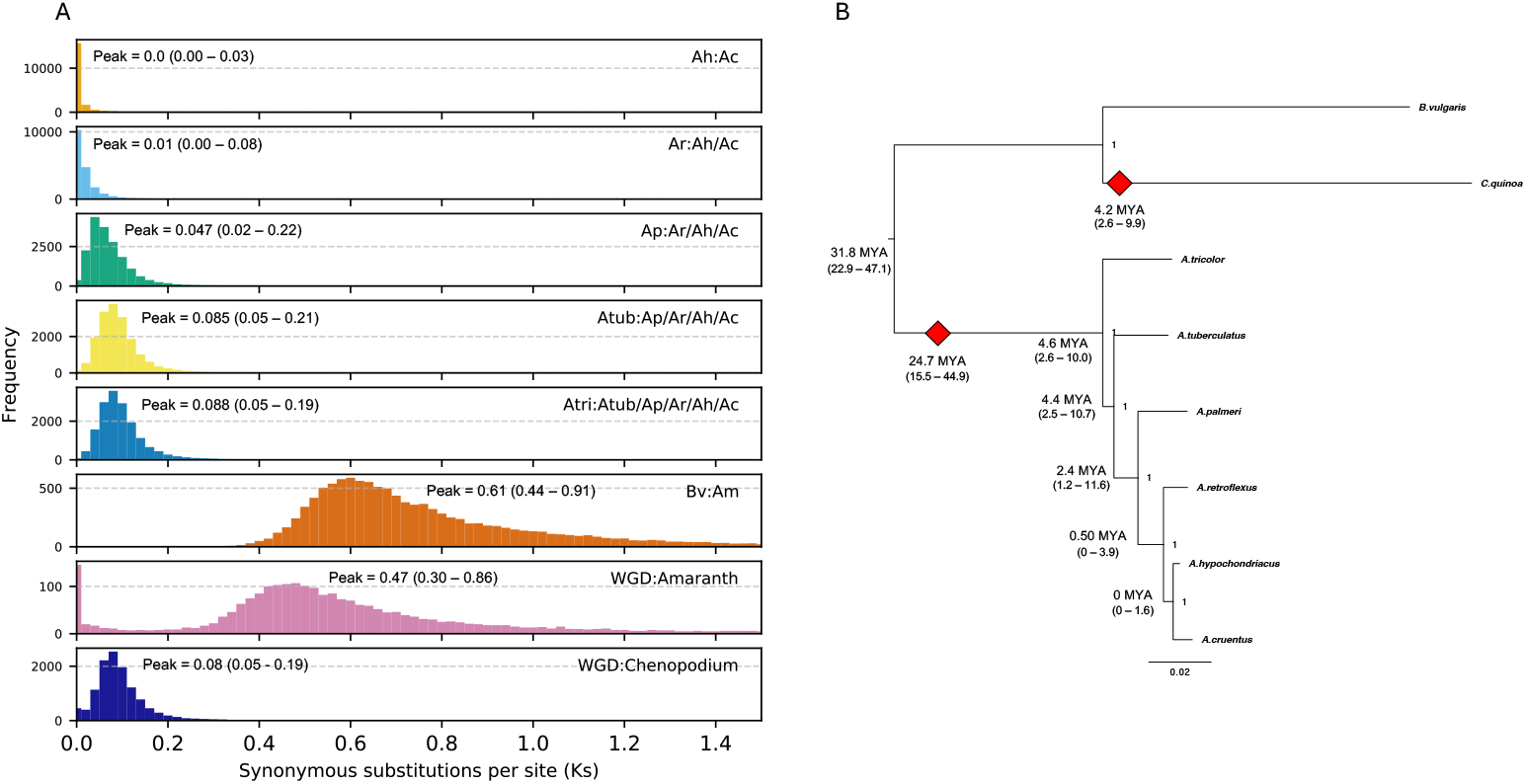
Synonymous substitutions per synonymous sites (K_s_) distribution identifying whole-genome duplication and speciation and events. **A** Synonymous substitutions per sites (Ks) histograms. Ks values of syntenic orthologs (speciation events) and paralogs (whole genome duplications) were determined using wgd2 [86]. Species abbreviations are *A. tricolor* (Atri), *A. tuberculatus* (Atub), *A. palmeri* (Ap), *A. retroflexus* (Ar), *A. cruentus* (Ac), *A. hypochondriacus* (Ah). Whole genome duplications (WGD) were determined by averaging across all *Amaranthus* species, while the WGD for *Chenopodium* was based solely on *Chenopodium quinoa*. Confidence intervals (95%) are provided in parenthesis. **B** Rooted *Amaranthus* species tree based on conserved orthologs as identified by Orthofinder2 [81] showing speciation and whole genome duplications dates as inferred by Ks distributions (see panel A). *B. vulgaris* (Bv) and *C. quinoa* were included as outgroups. Red diamonds indicate estimated points of WGD inferred from peaks in Ks.

**Figure E2.**
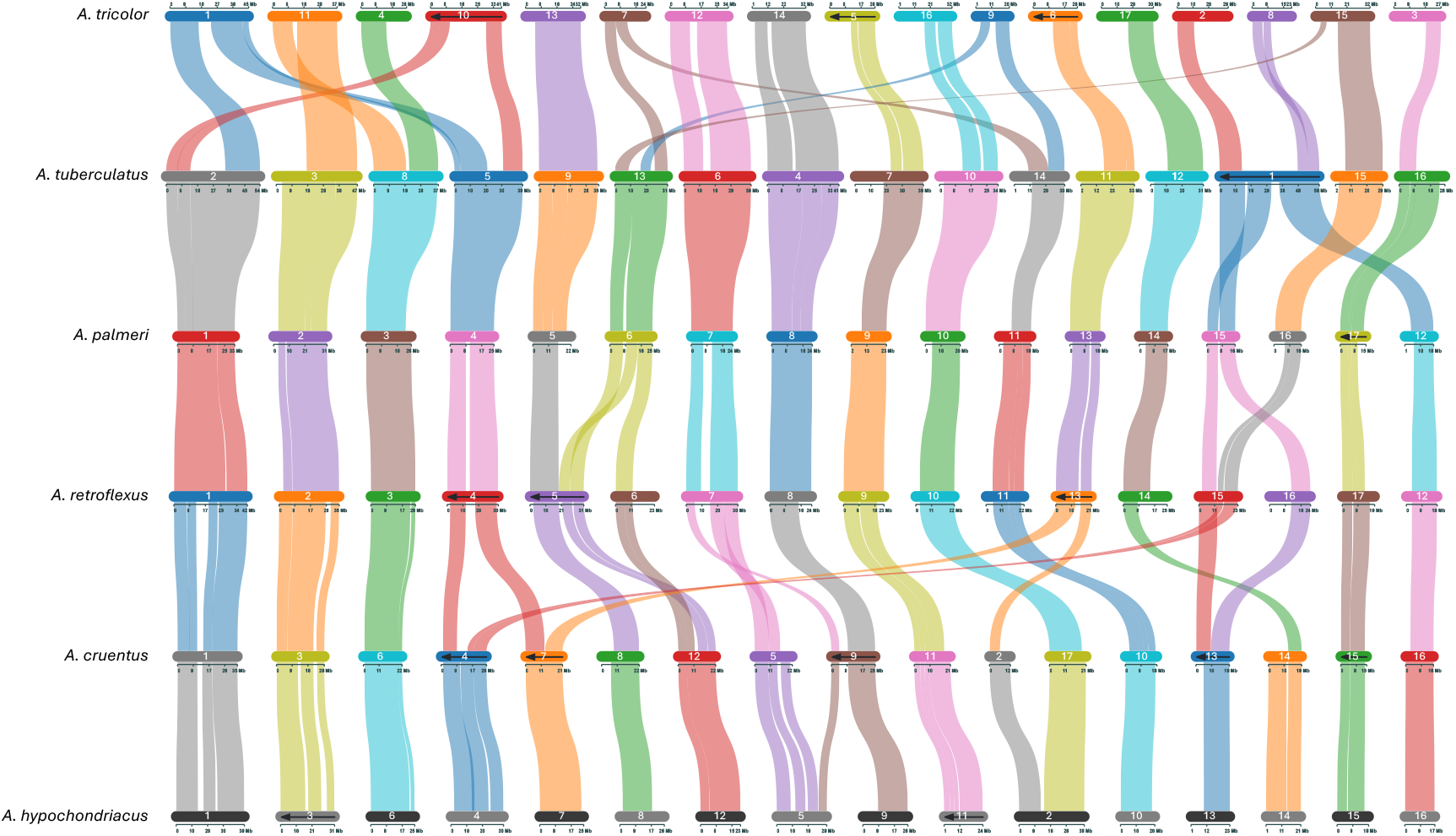
Syntenic relationships of orthologous regions among sequenced *Amaranthus* species. Species are ordered vertically by phylogenetic positions (*A. tricolor, A. tuberculatus, A. palmeri, A. retroflexus, A. cruentus* and *A. hypochondriacus*). Chromosomes are ordered horizontally, and ribbons are color-coded to show syntenic relationships among chromosomes. Arrows within chromosomes indicate reverse complementation.

**Figure E3.**
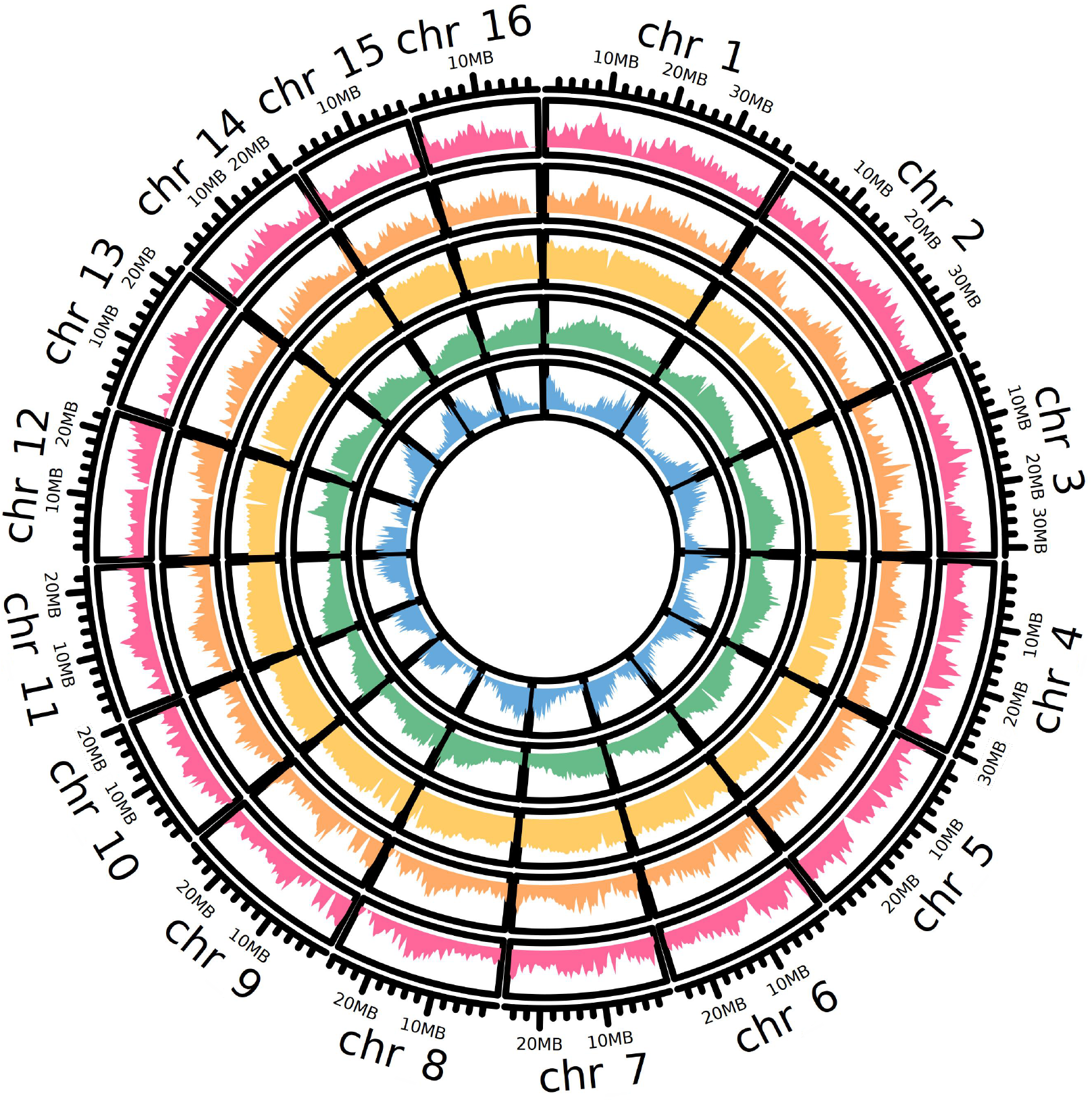
Genomic distribution of genes, methylation marks and accessible chromatin regions (ACRs) in *A. hypochondriacus* reference genome. Tracks depict the genomic distribution of (from inside to outside) annotated genes (blue), transposable elements (green), methylation in CpG context (yellow), ACRs found in leaf tissue (orange), ACRs found in seedling tissue (pink). The outer most ring represents the chromosomes of the reference genome and indicates the position within the chromosomes in megabase pairs (Mb).

**Figure E4.**
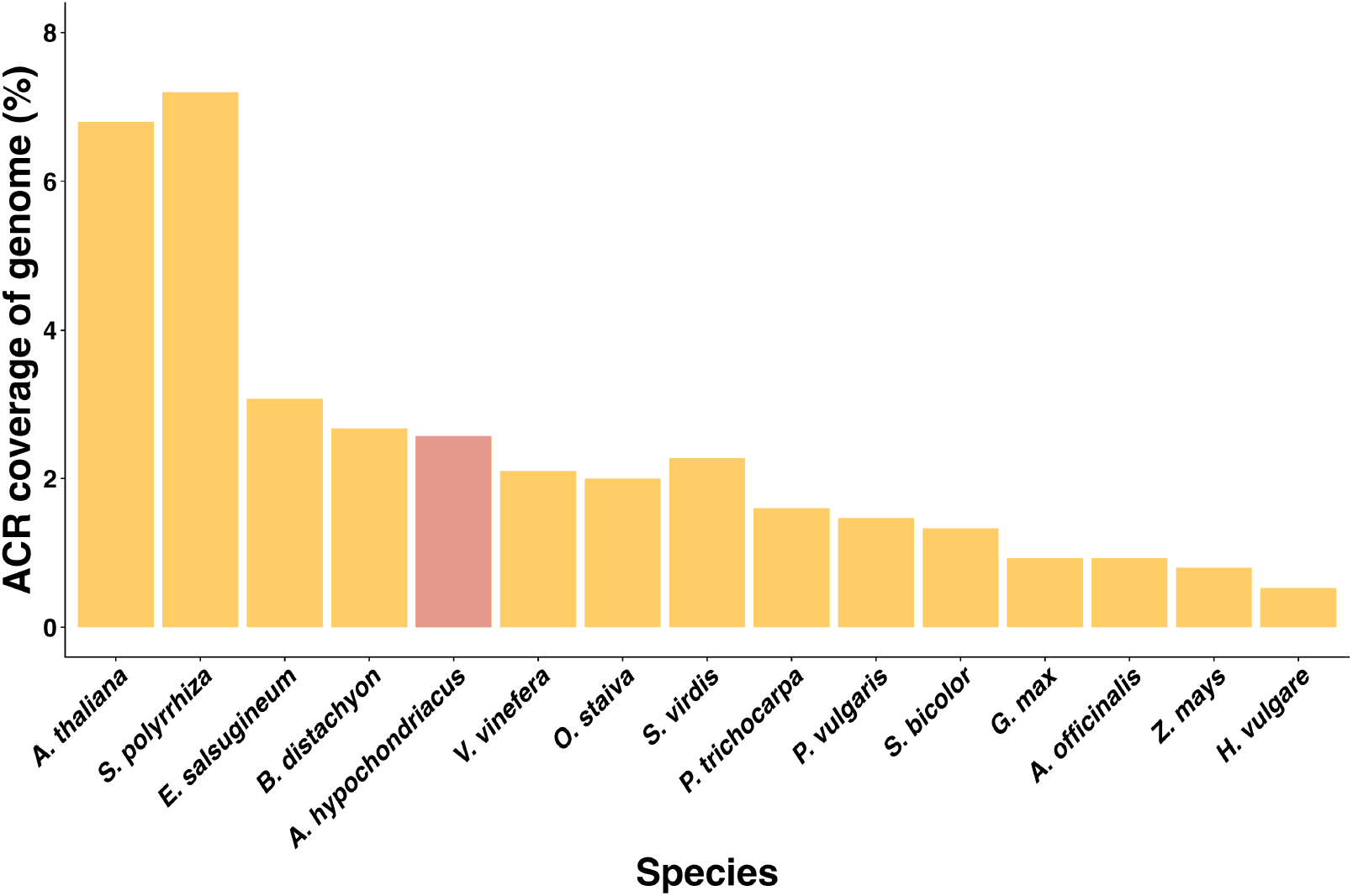
Fraction of accessible chromatin relative to genome size in 15 plant species. Y-axis indicates fraction of the whole genome (in %) that was found to be accessible. *A. hypochondriacus* (red) data based on mean coverage of PI 558499 samples, while the other species were plotted based on data from Lu et al. [35] and Schwope et al. [36]).

## Supplement

**Figure S1.**
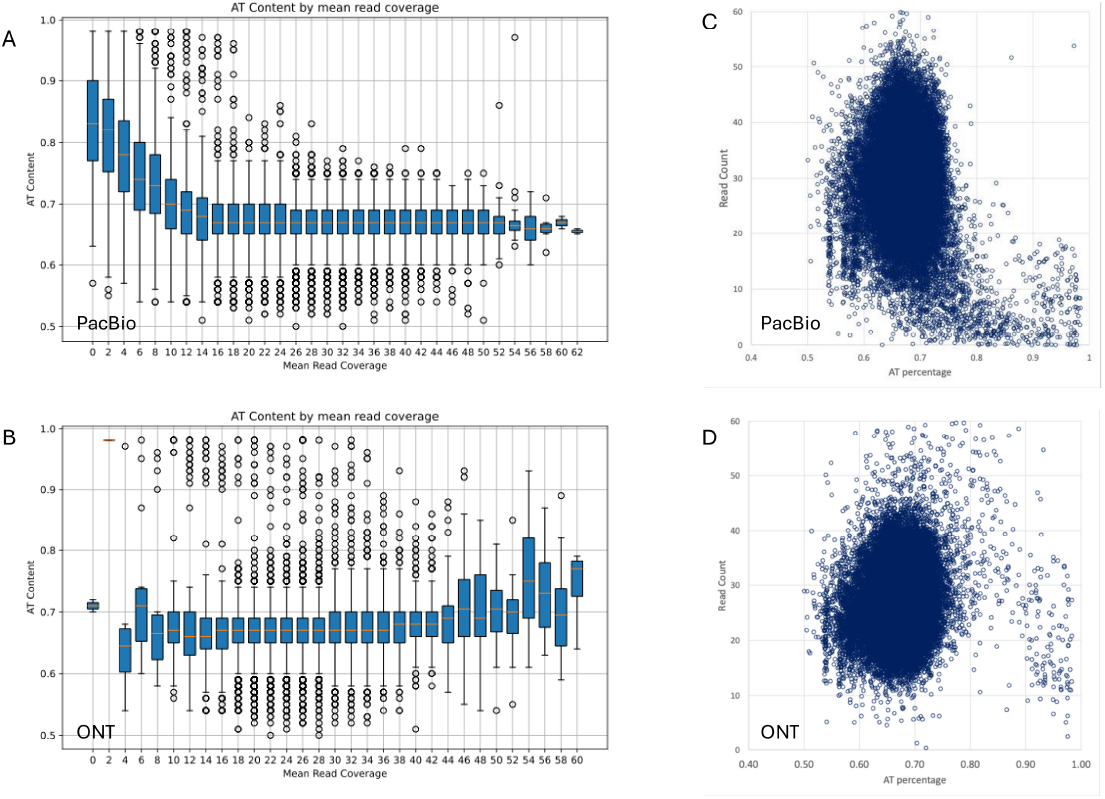
Read coverage relative to AT percentage with PacBio HiFi and ONT read types. Panels A (PacBio HiFi) and B (ONT) show the mean read depth (x-axis) at different AT percentage (y-axis) across 10 kb windows of the genome showing the AT bias of the PacBio reads, which leads to more fragmented primary assemblies. Similarly, panels C (PacBio HiFi) and D (ONT) show read count (y-axis) across 10 kb genomic windows with varying AT percentages (x-axis). Note the reduction of PacBio HiFi reads (panel C) in windows with high AT percentage. Amaranths are potentially unique in that they have regions of high AT concentration – at these regions, PacBio HiFi reads often terminate prematurely which then fragments the primary contig assembly. Using a hybrid approach (both PacBio and ONT) successfully resolves this issue.

**Figure S2.**
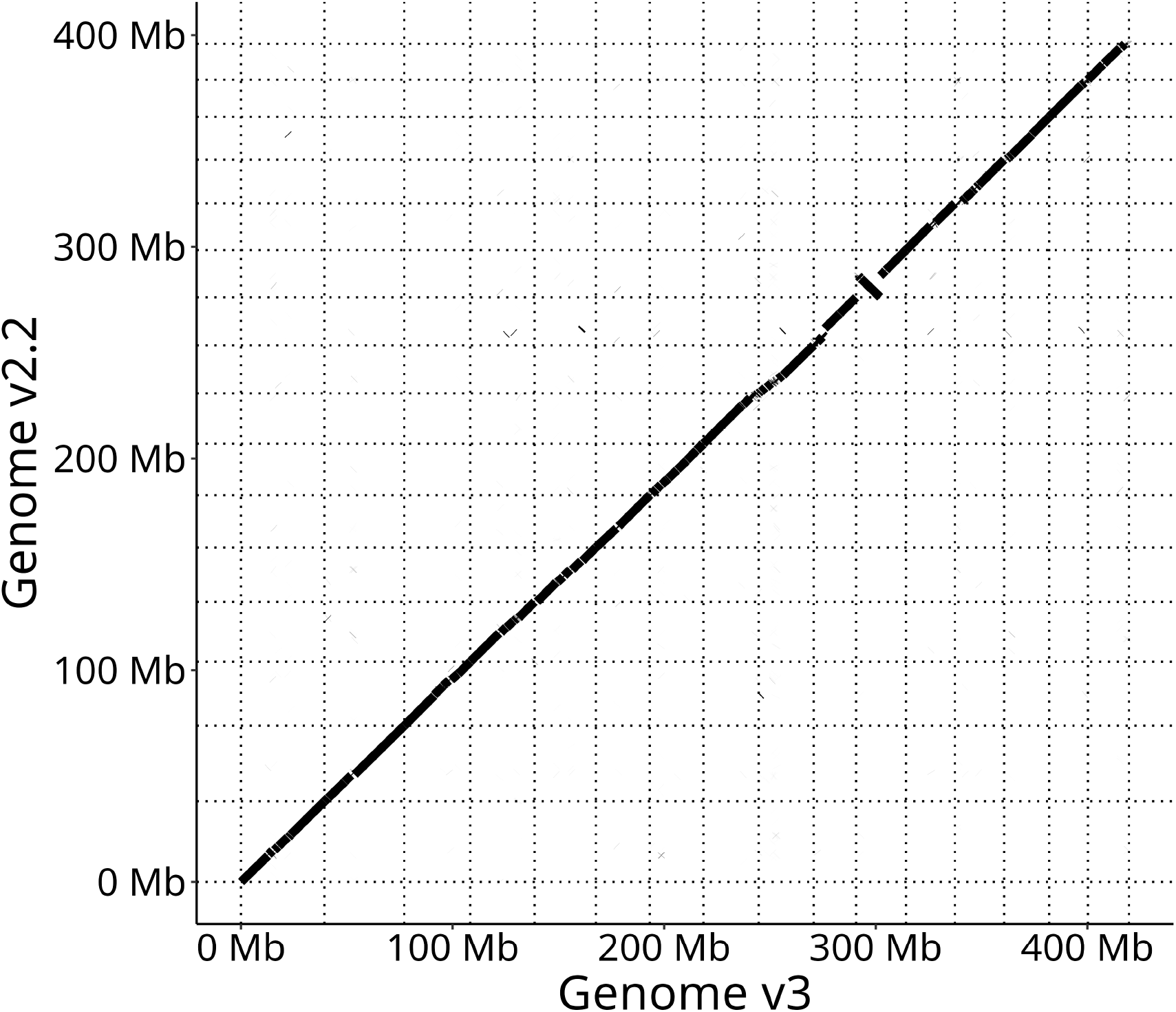
Dotplot of whole-genome alignment between *A. hypochondriacus* reference genome versions v2.2 and v3. Only scaffolds corresponding to the 16 chromosomes of *A. hypochondriacus* were depicted.

**Figure S3.**
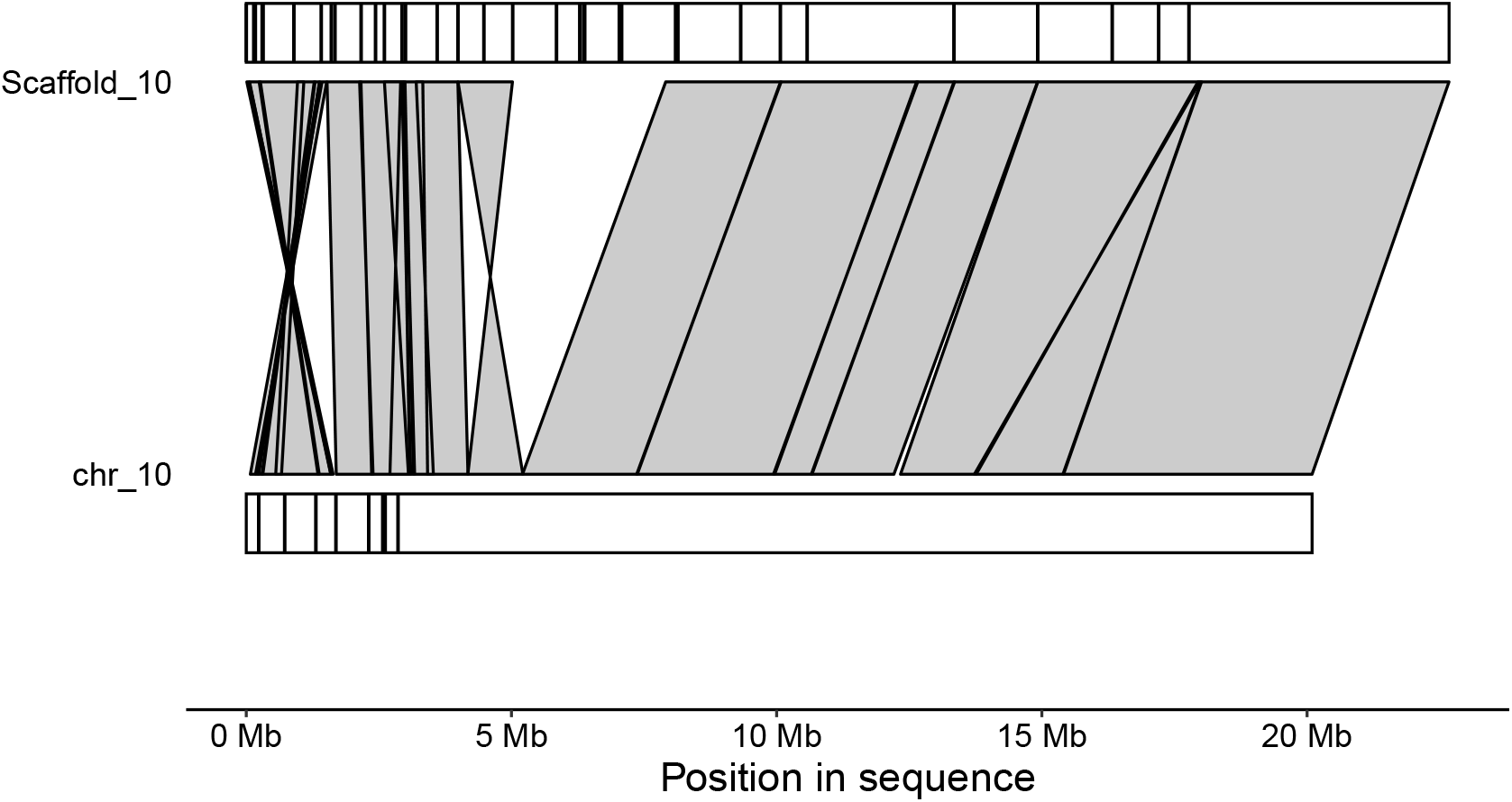
Synteny between *A. hypochondriacus* chromosome 10 scaffolds of reference genome versions v2.2 and v3. Alignment of scaffold 10 of reference assembly v2.2 and chromosome 10 of reference assembly v3 shows potentially misassembled region missing from the new chromosome assembly. Contig borders of assembly v2.2 and v3 are indicated as vertical black bars on their respective chromosomes.

**Figure S4.**
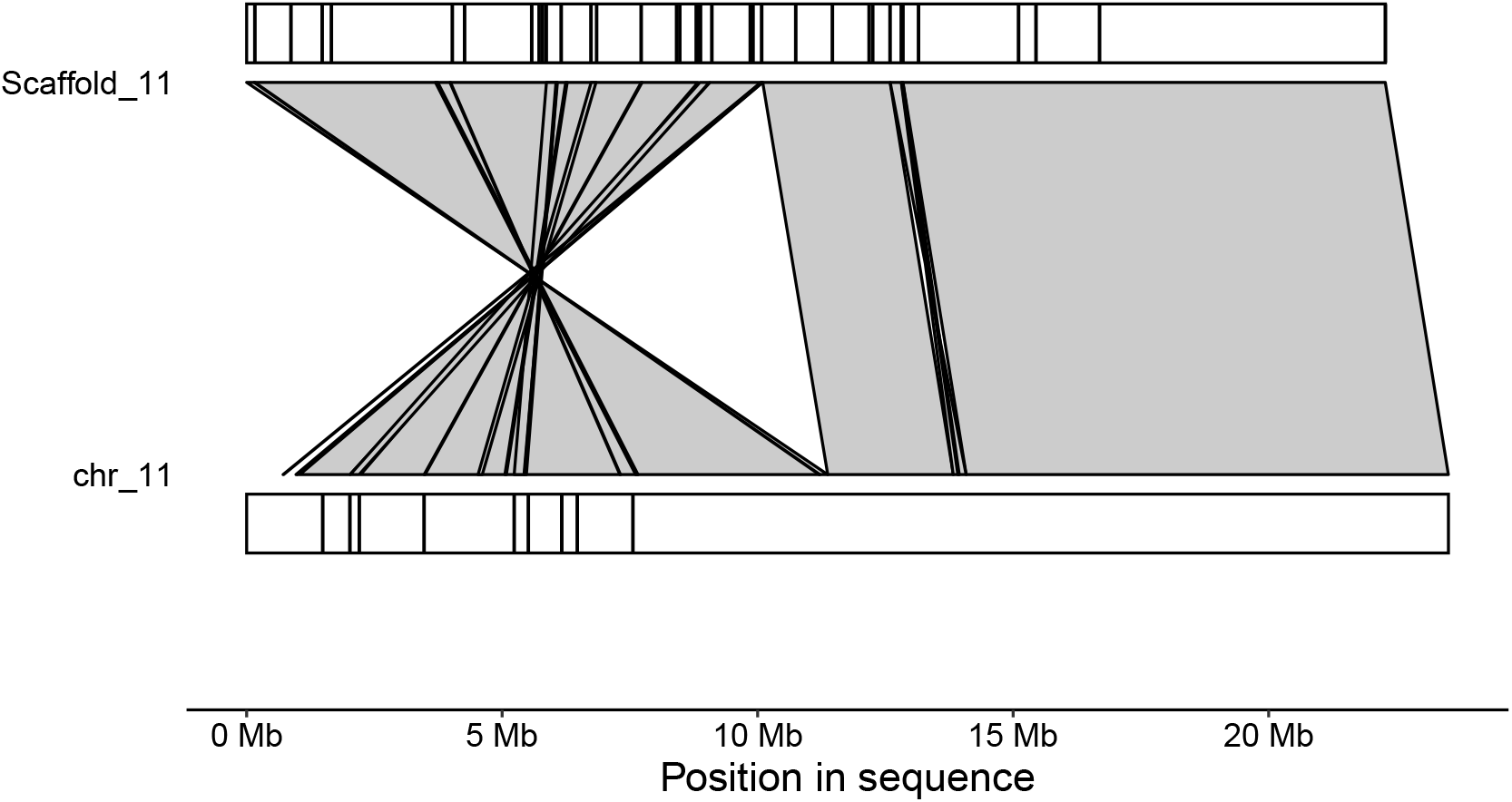
Synteny between *A. hypochondriacus* chromosome 11 scaffolds of reference genome versions v2.2 and v3. Alignment of scaffold 11 of reference assembly v2.2 and chromosome 11 of reference assembly v3 shows potentially misassembled region inverted between the two assemblies. Contig borders of assembly v2.2 and v3 are indicated as vertical black bars on their respective chromosomes.

**Figure S5.**
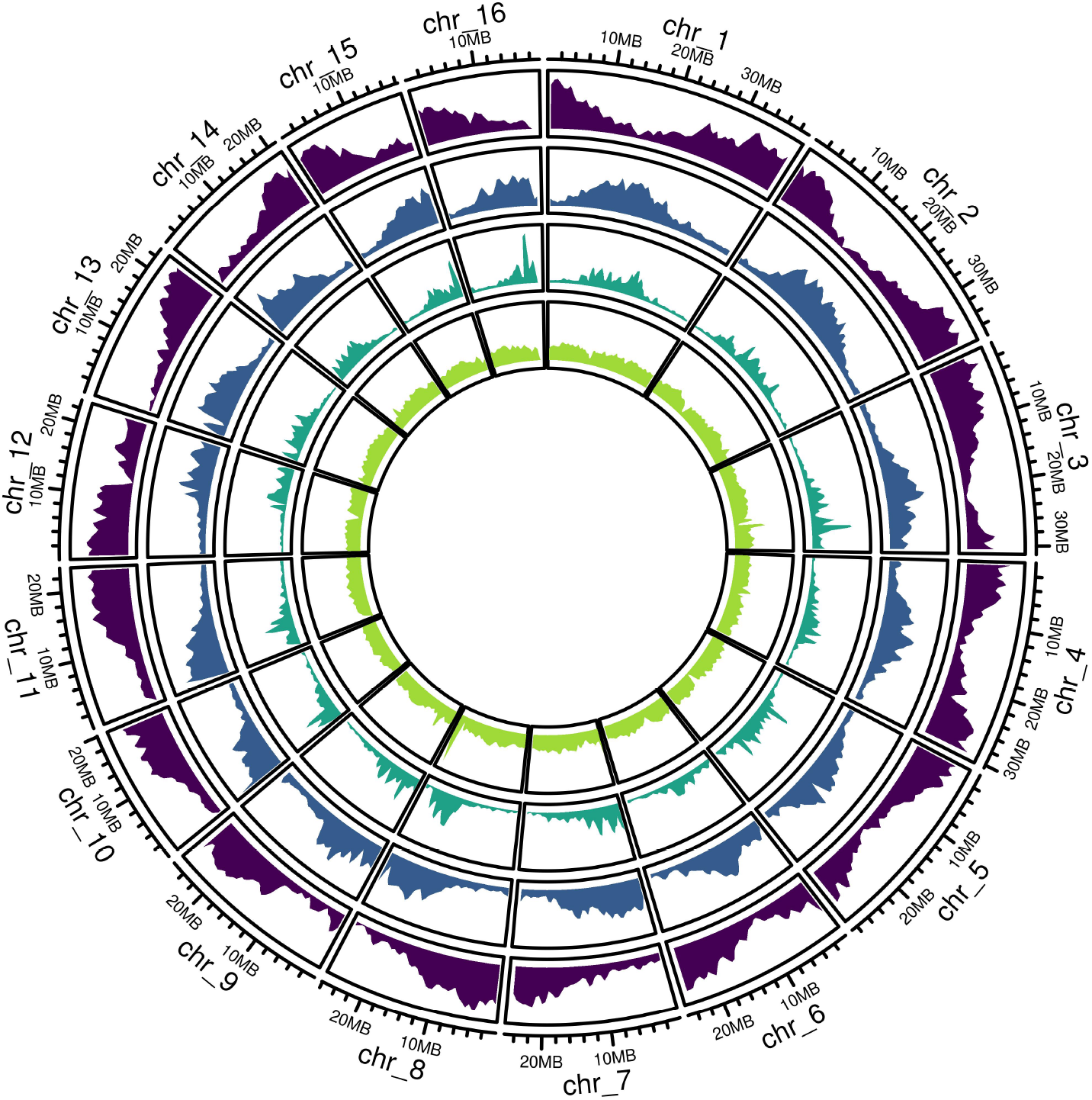
Genomic distribution of annotated genes and transposable element classes in *A. hypochondriacus* reference genome v3. Tracks depict (from outside to inside) density of genes, LTR elements, LINEs, and MITEs calculated in 1 Mb windows along the genome.

**Figure S6.**
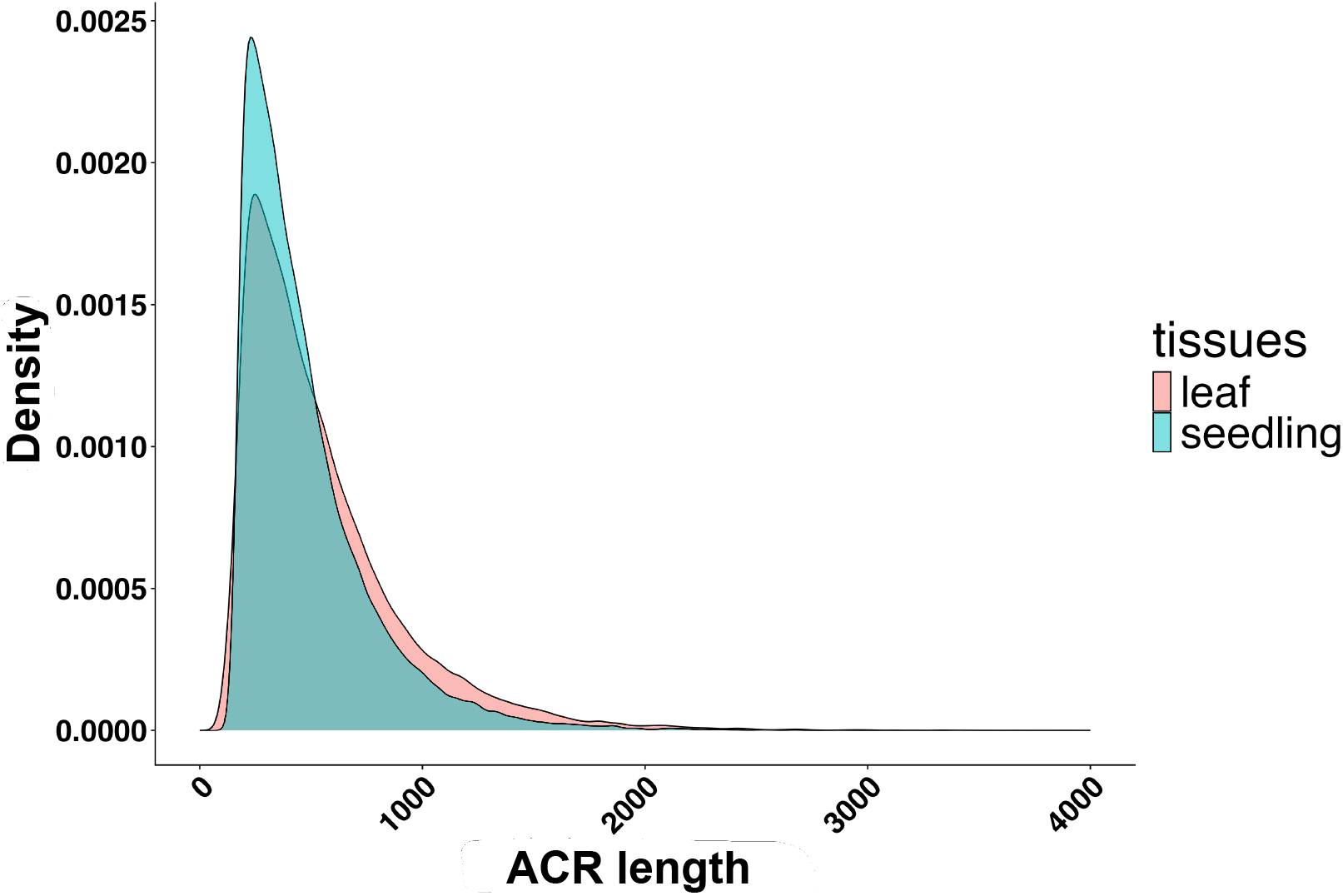
Distribution of ACR length in the ATACome. Length of both leaf tissues (pink) and seedling tissue (turquoise) ACRs from samples of PI 558499 aligned to the *A. hypochondriacus* reference genome.

**Figure S7.**
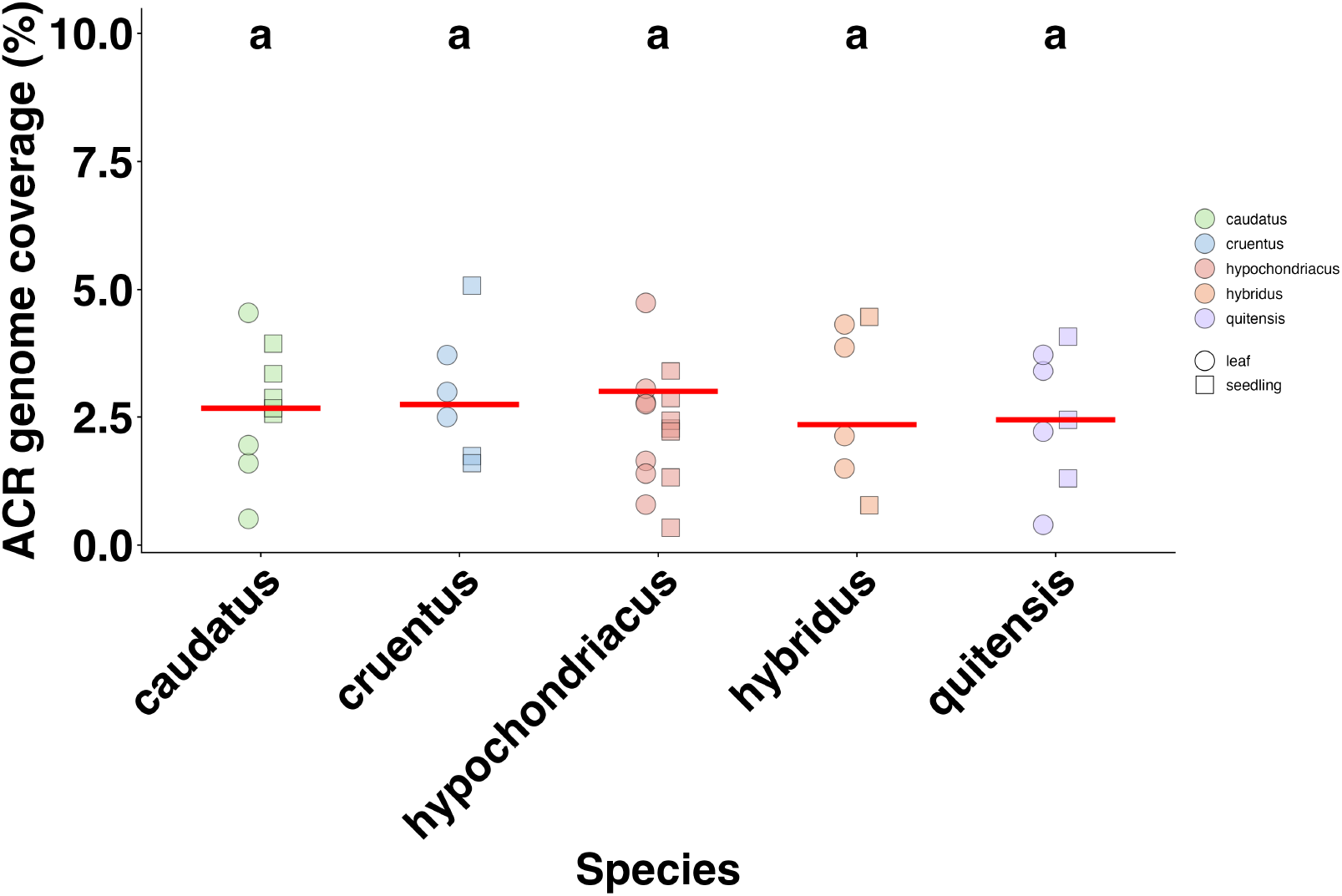
Fraction of the *A. hypochondriacus* reference genome covered by the ACRs called in each sample. The tissue and species from which each sample originates is indicated by shape and color, respectively. The mean is indicated by a red line for each species, respectively. Species for which the mean does not significantly differ are indicated by the same letter atop their column.

**Figure S8.**
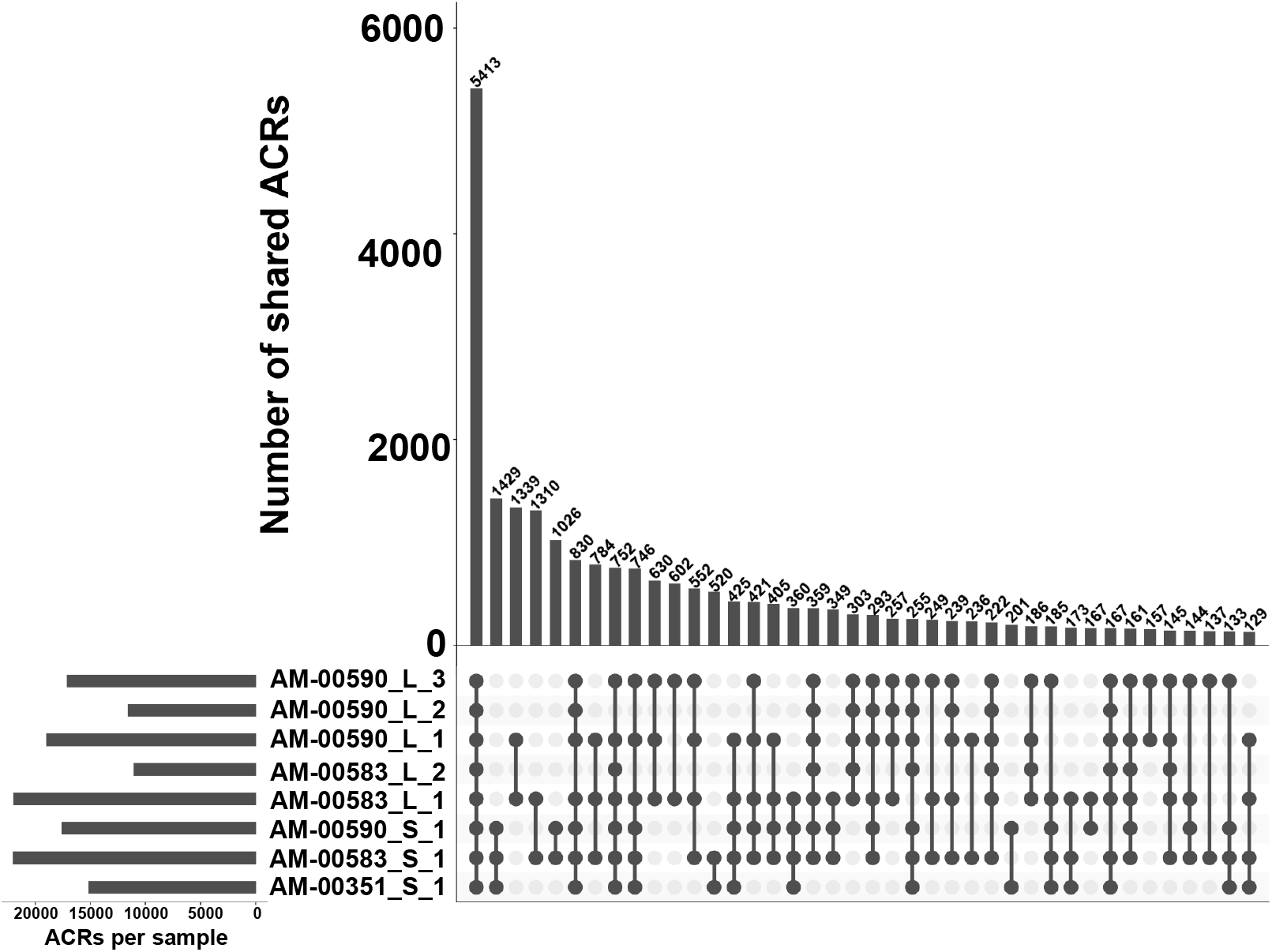
Overlap of ACRs between the samples of *A. hypochondriacus*. UpsetR plot of ACRs called in each of the eight samples from PI 558499. The Y-axis of the bar graph indicates the number of ACRs that were shared by samples indicated by joint dots in the corresponding column of the matrix below. ACRs are only part of one column. The second bar graph to the left of the sample names indicates total number of ACRs called in each sample.

**Figure S9.**
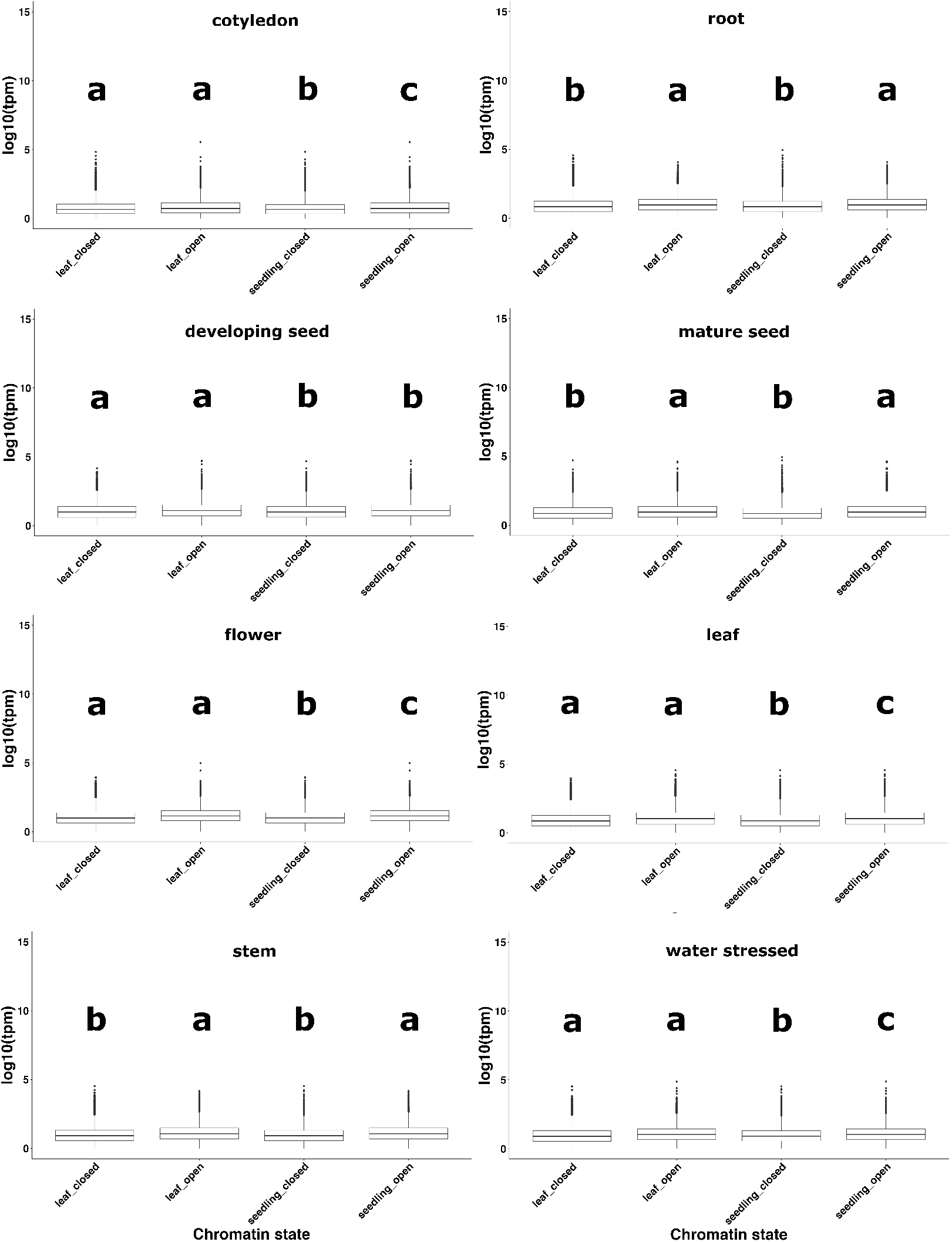
Expression comparison in eight tissues. Comparison of expression of genes associated with accessible and closed chromatin in leaf and seedling samples. Genes including promoters (2 kb upstream) that overlapped with at least one ACR were defined as open. An equally sized random subsample of ‘closed genes’ was taken as comparison. Mean expression values for the genes in each of the eight tissues was taken from [71]. Letters on top of the boxplots indicate significant difference within each of the eight tissues.

**Figure S10.**
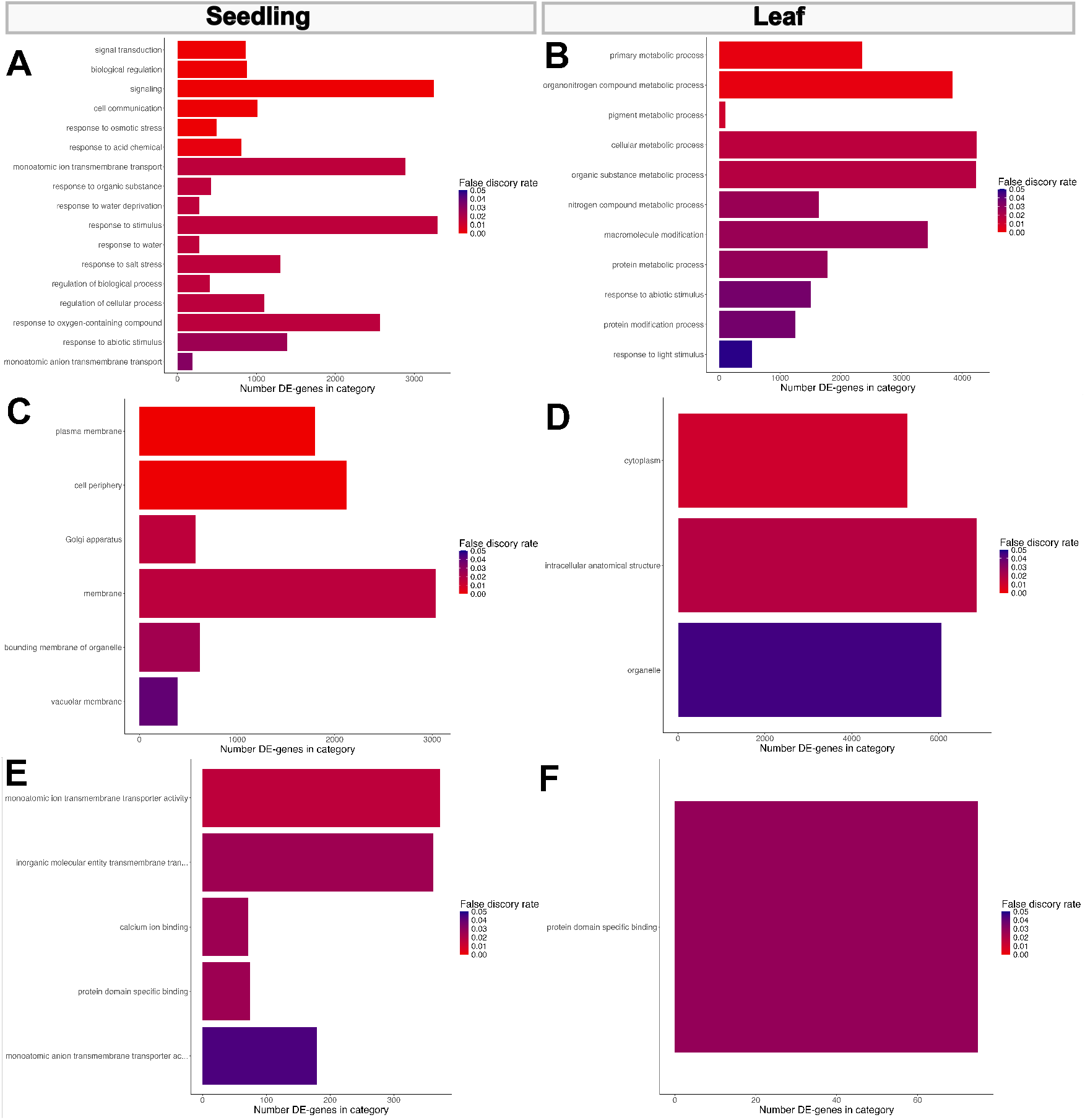
Functional categories enriched among genes associated with ACRs in *A. hypochondriacus*. GO-terms enriched among genes associated with ACRs for **A**. biological processes in seedling tissue **B**. biological processes in leaf tissue **C**. cellular components in seedling tissue **D**. cellular components in leaf tissue and **E**. molecular functions in seedling tissue **F**. molecular functions in leaf tissue. False discovery rate cutoff was set 0.05.

**Figure S11.**
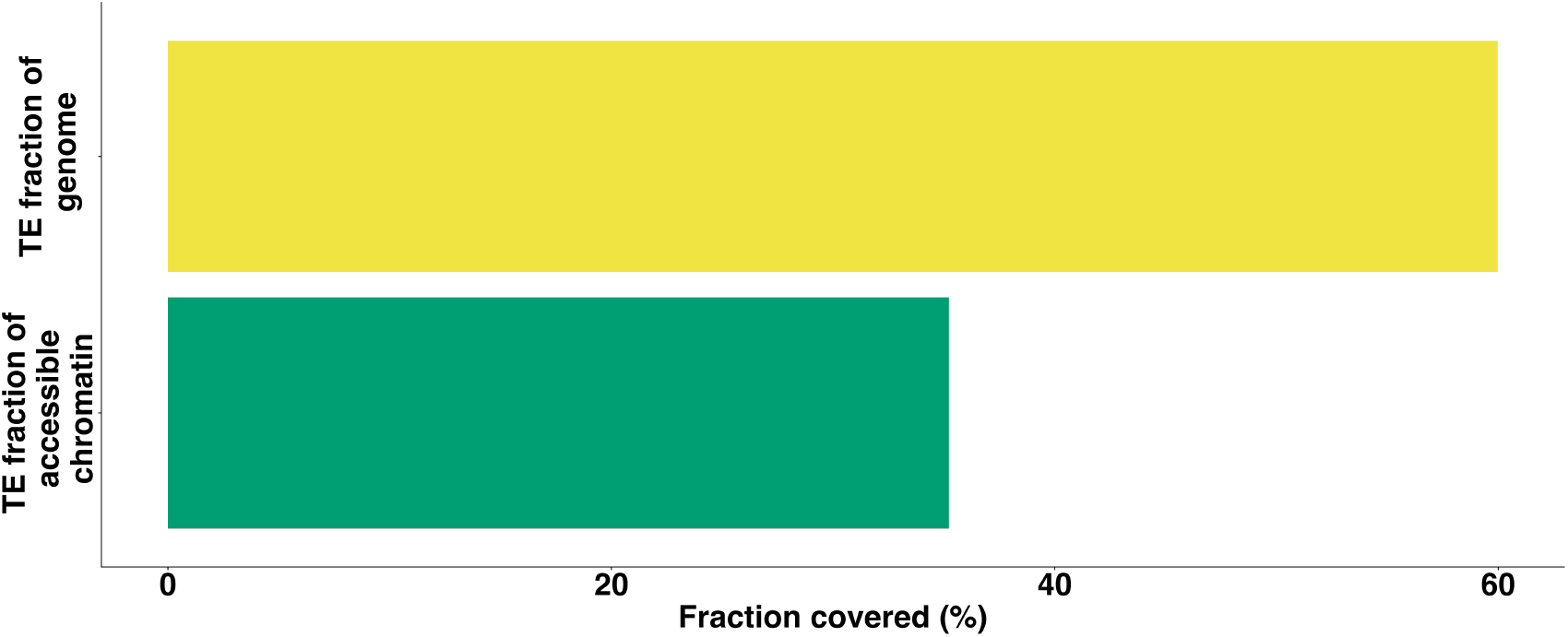
TE coverage of open chromatin and the genome. X-axis indicates the fraction of the whole genome (yellow) of *A. hypochondriacus* and accessible chromatin (green) that is occupied by TEs.

**Figure S12.**
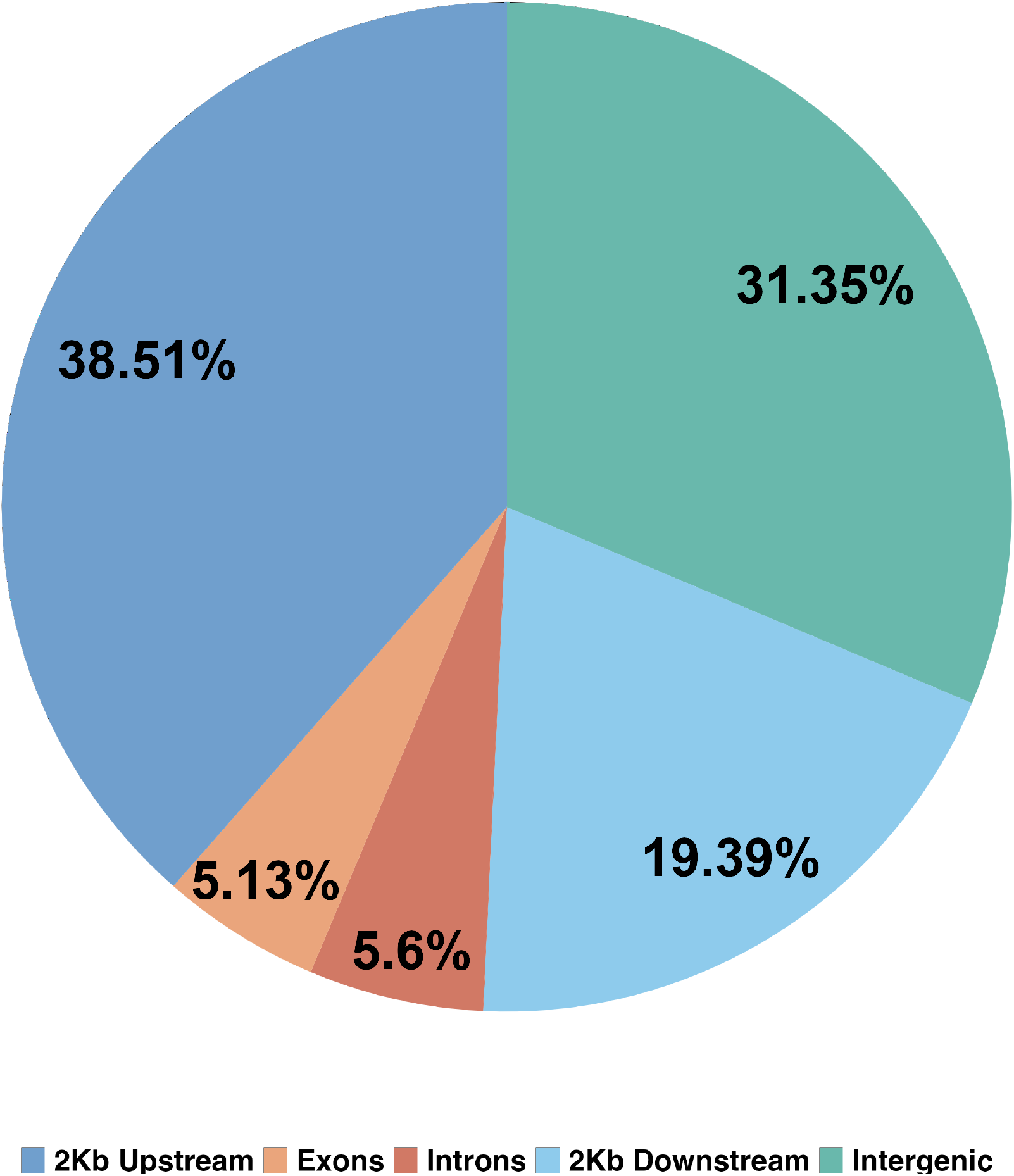
ACR distribution among the genomic regions in *A. cruentus*. ACRs were called from eight *A. hypochondriacus* samples of the reference accession PI 558499 aligned to the *A. cruentus* reference genome to test for reference bias. The genome was split into 8 categories, i.e., 2 kb upstream of transcription start site (TSS), Exons, Introns, 2 kb downstream of transcription termination site (TTS) and intergenic (everything not within 2 kb of TSS or TTS of a gene)

**Figure S13.**
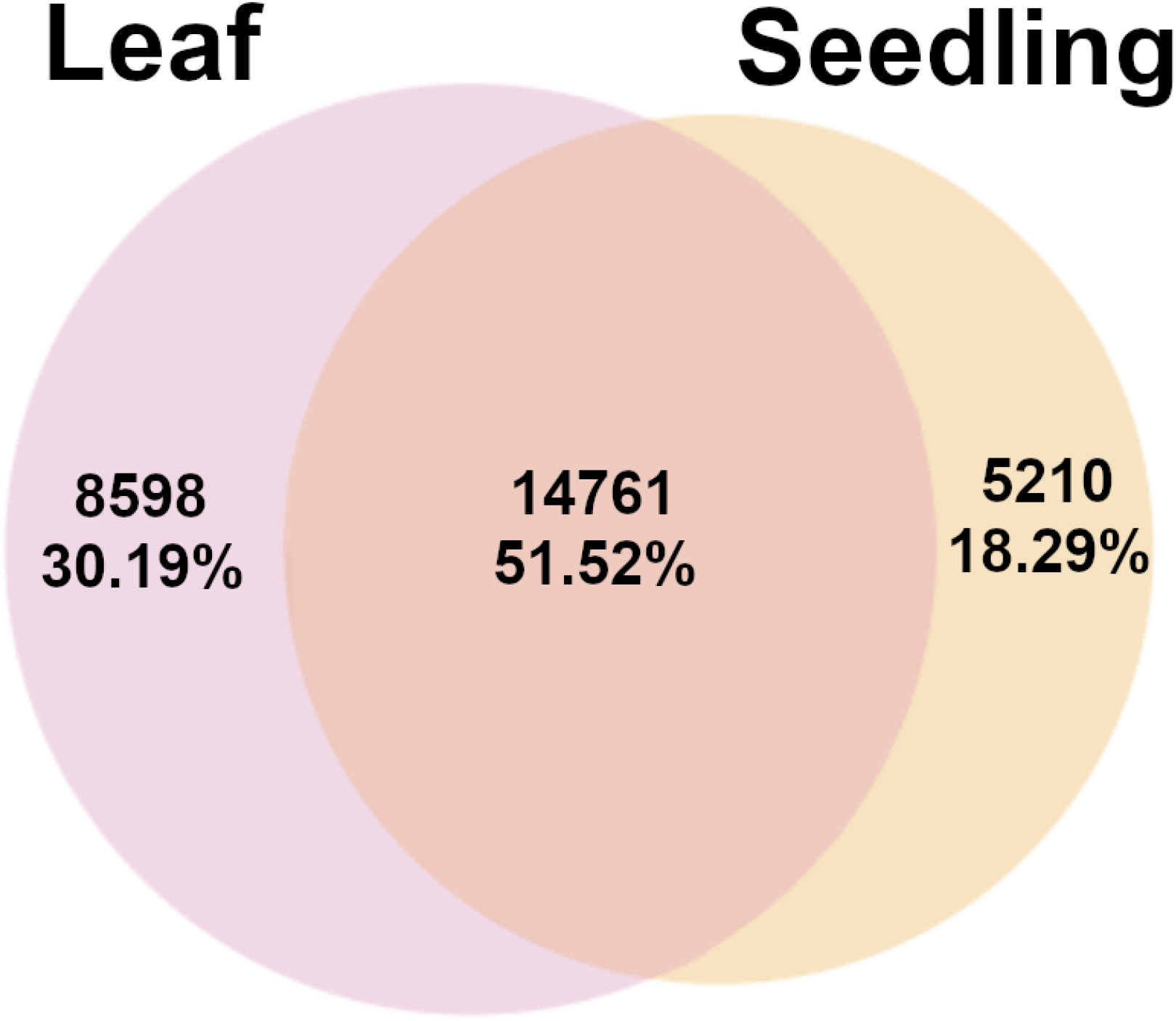
Overlap of ACRs between tissues in *A. cruentus*. ACRs called from eight samples of the *A. hypochondriacus* reference accession PI 558499 aligned to the *A. cruentus* reference genome to test for reference bias. Only ACRs that occurred in at least two samples of each tissue respectively were considered. ACRs from two samples of the same tissue that overlapped were joined together. ACRs had to overlap by at least 1 bp to be considered shared between the tissues

**Figure S14.**
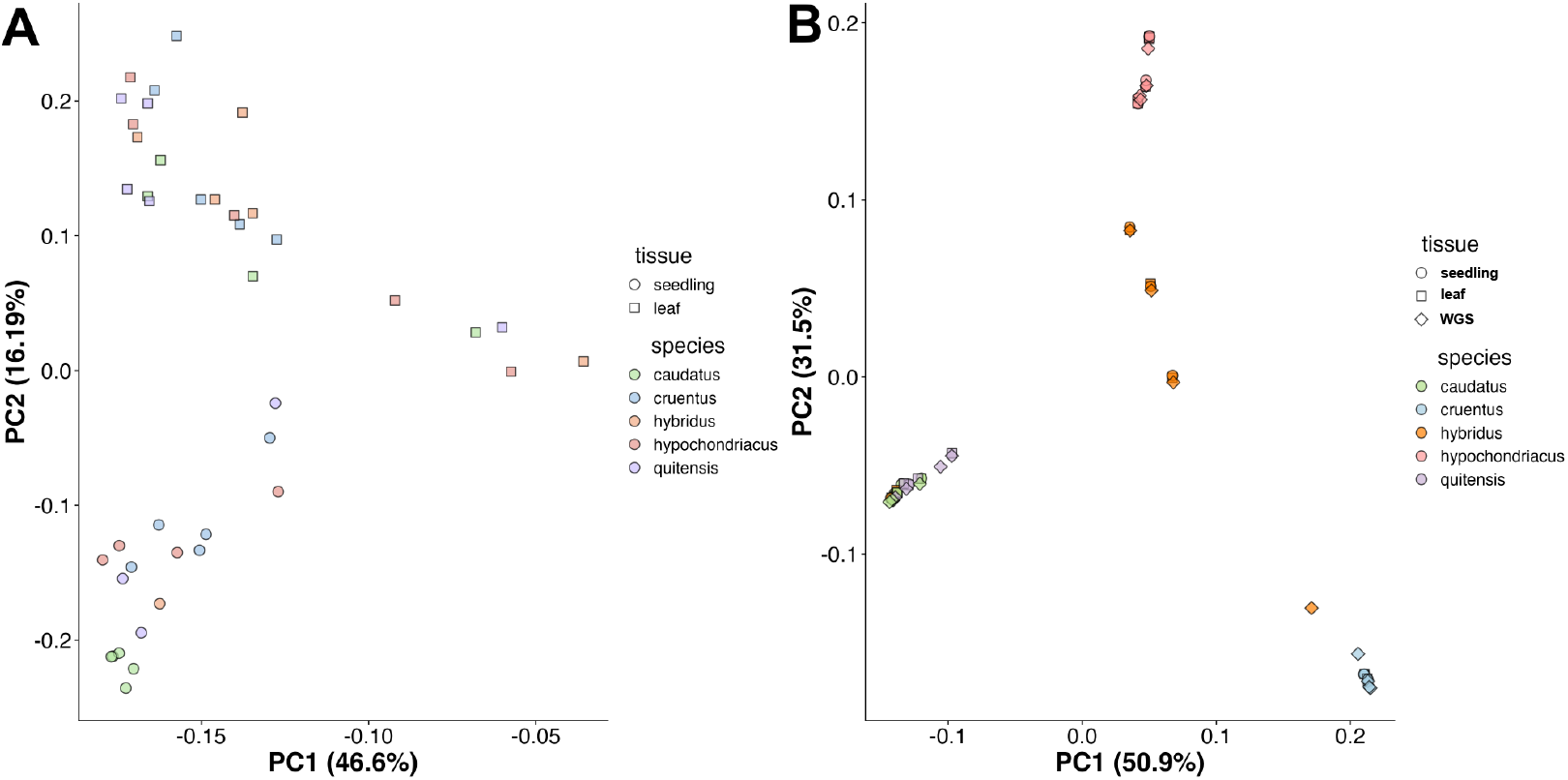
Relationship between ATAC-samples based on accessible chromatin regions (ACRs) and genome-wide SNPs. PCA of the 42 ATAC-seq samples based on **A**. the 51,571 ACRs called across samples. The sampled tissue and species is indicated by shape and color, respectively. **B**. PCA based on SNP data from ATAC-sequencing data and whole genome sequencing data (WGS) showing that ATAC-seq data recovers the population structure.

**Figure S15.**
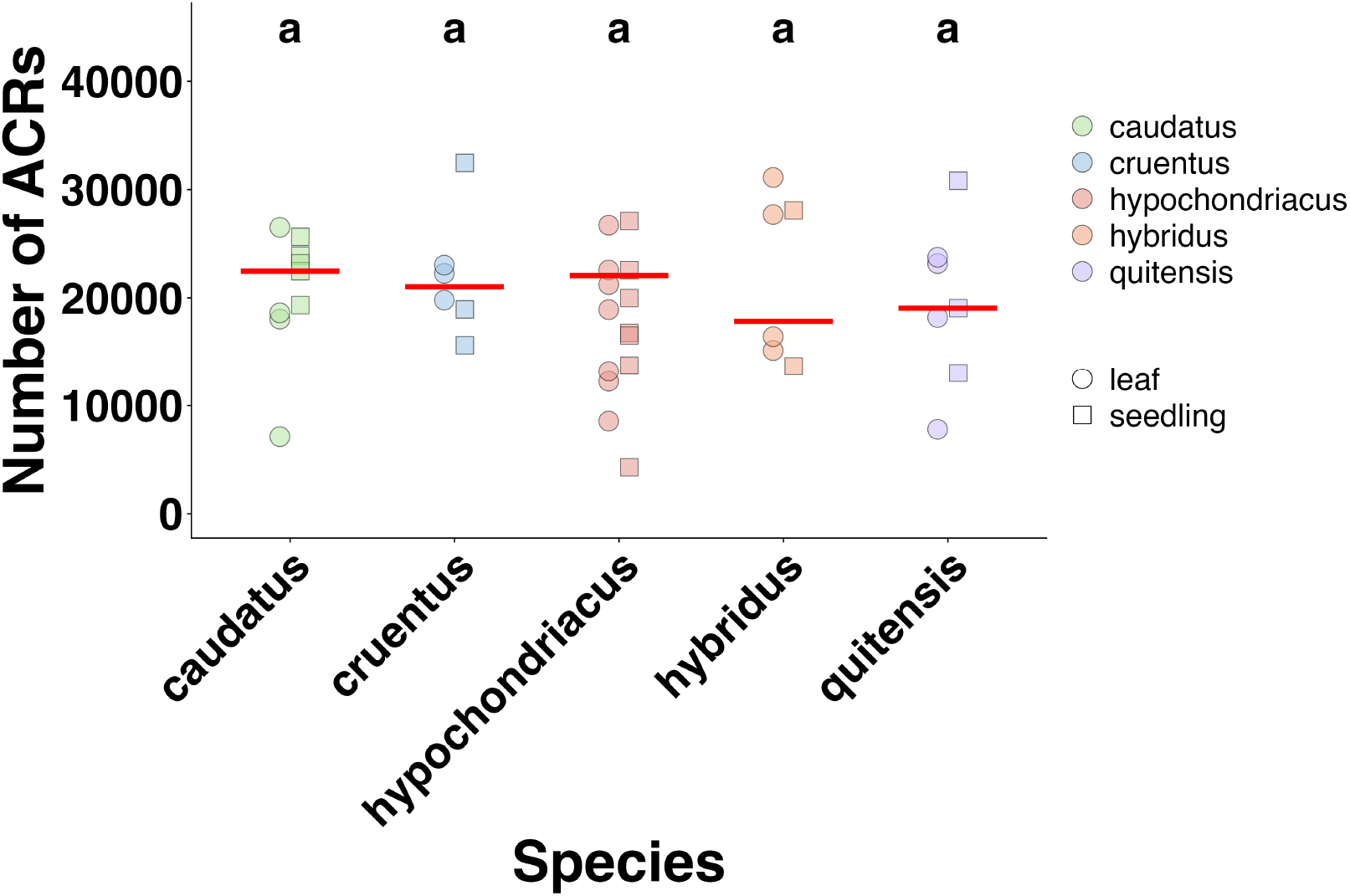
Number of ACRs called for each sample when aligned to the *A. hypochondriacus* reference genome. The tissue and species from which each sample originates is indicated by shape and color, respectively. The mean is indicated by a red line for each species, respectively. Species for which the mean does not significantly differ are indicated by the same letter atop their column.

**Figure S16.**
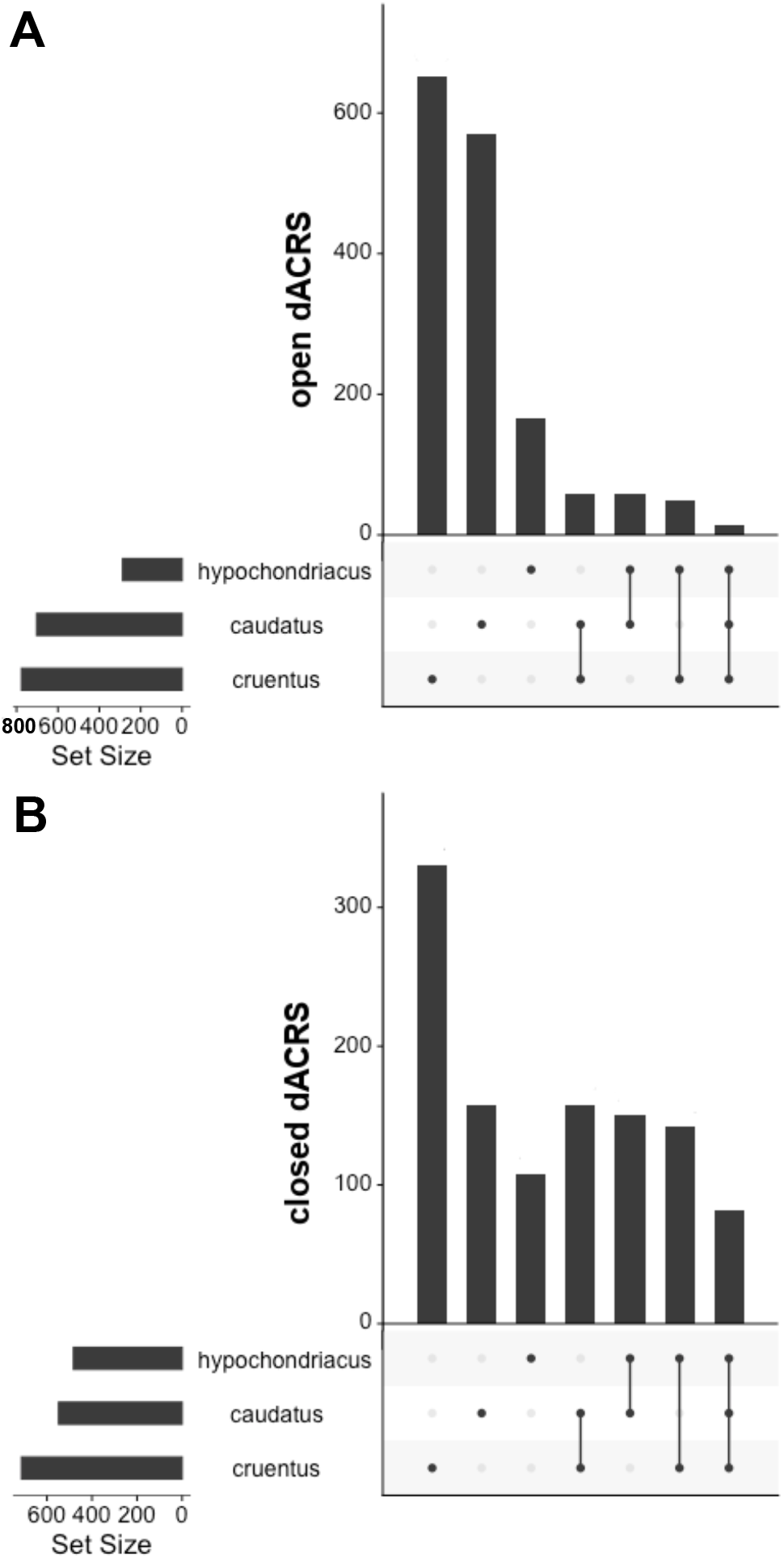
Number of dACRs that were either unique to each domesticate or shared between them. UpsetR plot of ACRs that are differentially accessible (dACRs) between the domesticates and their wild ancestor. **A**. dACRs that opened during domestication. **B**. dACRs that closed during domestication.

**Figure S17.**
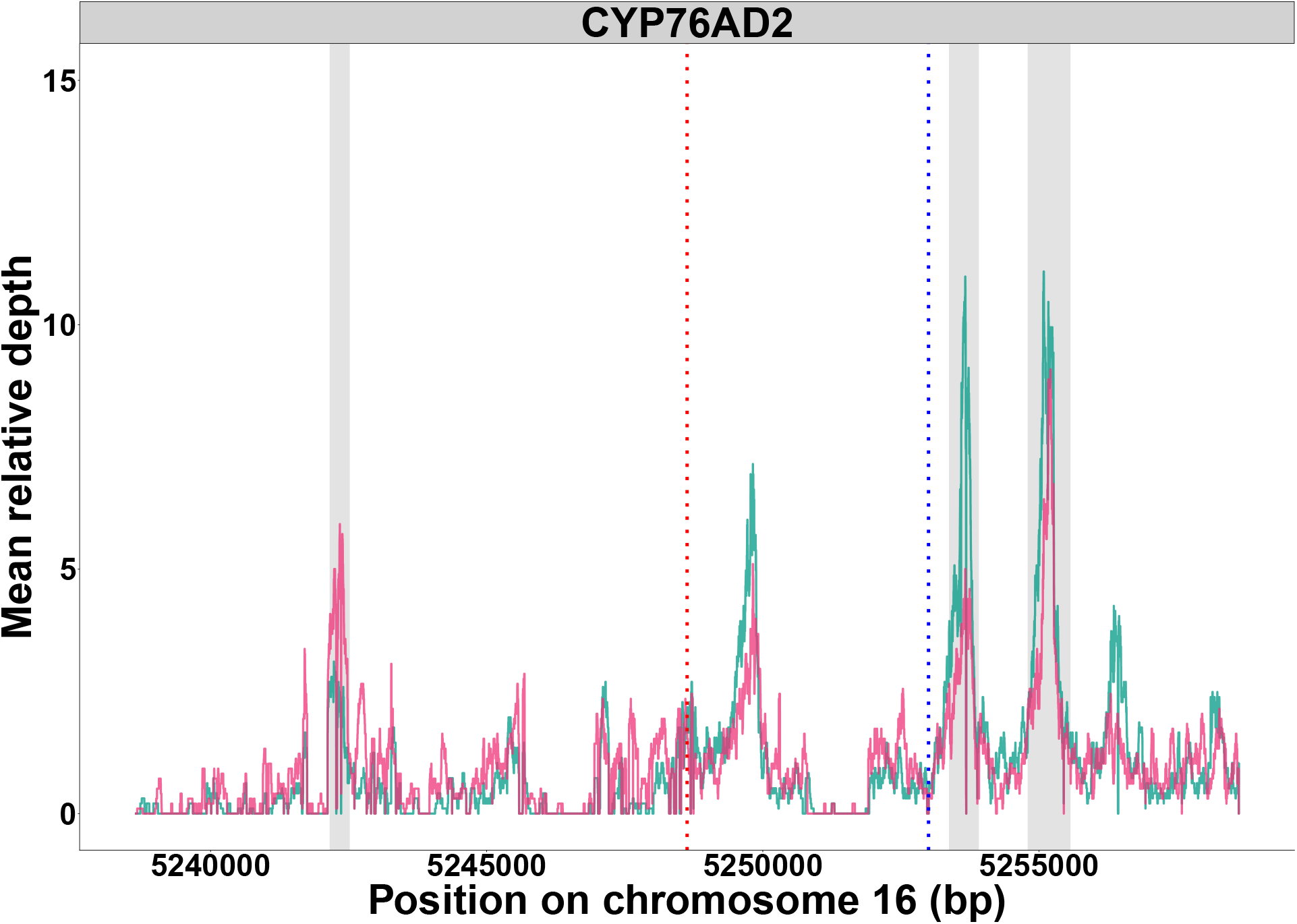
**Mean normalized read depth 10 kb up and downstream of the transcription start site (TSS) of CYP76AD2 for *A. caudatus* Accessions with red and green seedling color (magenta and green respectively). Vertical dotted lines indicate TSS (red) and Transcription End Site (TES) (blue). Grey vertical areas indicated called accessible chromatin regions (ACRs). Normalized depth was calculated by determining the mean depth for each site across accessions with red and green seedlings respectively and normalized by the mean depth of the whole region**.

## Supplementary Tables

**Table S1.**
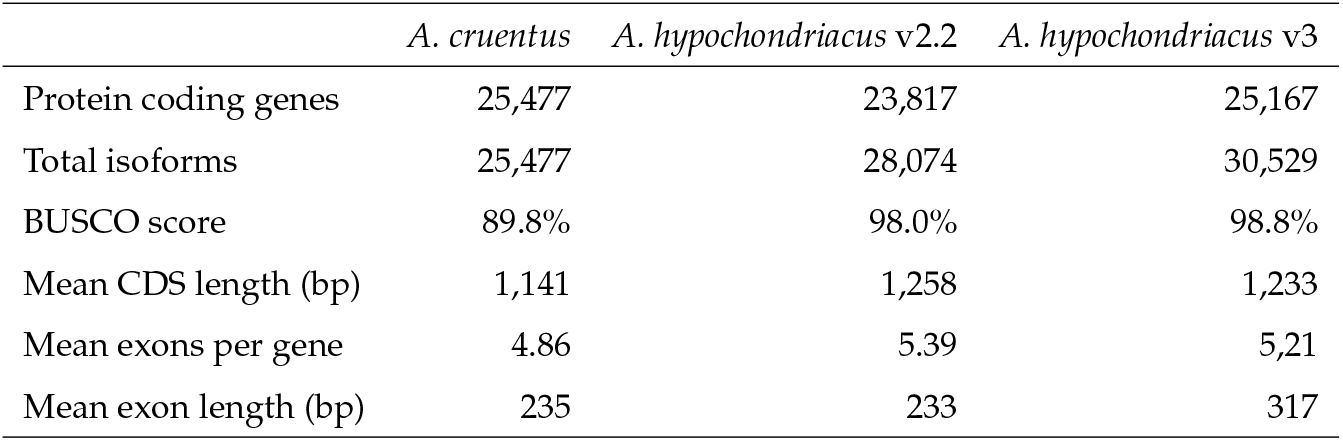
Comparison of genome annotation statistics between published grain amaranth reference genomes.

**Table S2.**
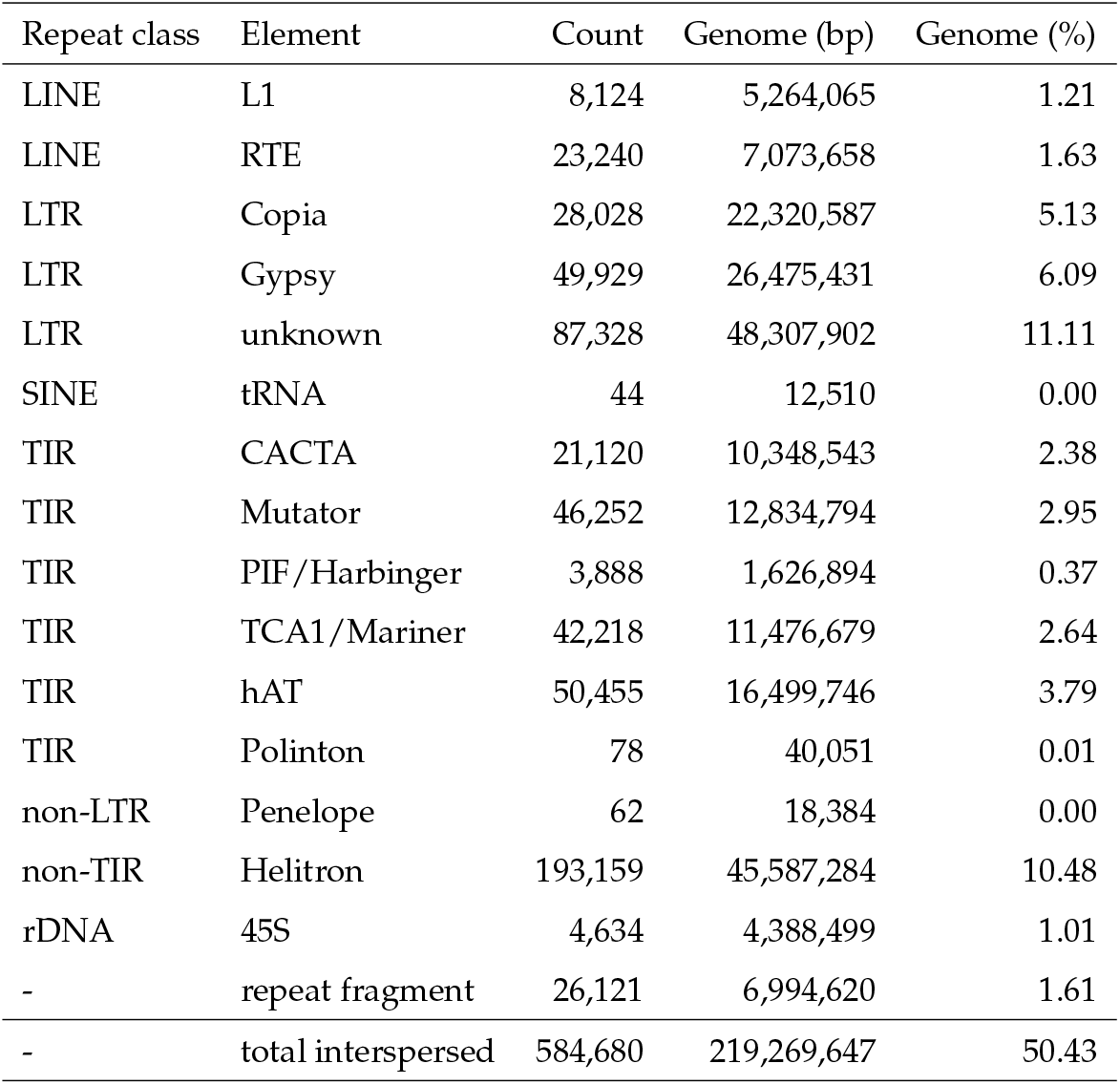
Transposable element annotation summary statistics for the *A. hypochondriacus* genome annotation v3.0. Displayed are the repetitive element class (if applicable) and subclass, the number of elements identified and the fraction of the genome they make up.

**Table S3.** Overview of samples used in this study.

| Accession ID | Species | Sample | Mapped Reads | Called Peaks |
| --- | --- | --- | --- | --- |
| PI 490518 | *A. caudatus* | seedling | 26324188 | 25478 |
| PI 490518 | *A. caudatus* | leaf | 44428081 | 37937 |
| PI 490518 | *A. caudatus* | leaf | 44672690 | 31659 |
| PI 490612 | *A. caudatus* | seedling | 37597522 | 36760 |
| PI 490612 | *A. caudatus* | leaf | 24522965 | 18941 |
| PI 490612 | *A. caudatus* | seedling | 51173694 | 45953 |
| PI 608019 | *A. caudatus* | leaf | 40953093 | 39437 |
| PI 608019 | *A. caudatus* | seedling | 42451784 | 37165 |
| PI 642741 | *A. caudatus* | seedling | 28311405 | 31401 |
| PI 642741 | *A. caudatus* | seedling | 37197967 | 30450 |
| PI 642741 | *A. caudatus* | leaf | 19980225 | 27455 |
| PI 642741 | *A. caudatus* | leaf | 19404739 | 18318 |
| PI 511714 | *A. cruentus* | leaf | 48802567 | 43012 |
| PI 511714 | *A. cruentus* | seedling | 45235760 | 44699 |
| PI 511717 | *A. cruentus* | leaf | 35981869 | 33253 |
| PI 511717 | *A. cruentus* | seedling | 29246205 | 26832 |
| PI 643058 | *A. cruentus* | leaf | 46230710 | 39692 |
| PI 643058 | *A. cruentus* | seedling | 22968690 | 25020 |
| PI 490489 | *A. hybridus* | leaf | 34286717 | 40907 |
| PI 490489 | *A. hybridus* | seedling | 41484966 | 43143 |
| PI 511754 | *A. hybridus* | leaf | 10576068 | 24806 |
| PI 511754 | *A. hybridus* | leaf | 34048324 | 28814 |
| PI 652426 | *A. hybridus* | seedling | 42521074 | 38959 |
| PI 558499 | *A. hypochondriacus* | seedling | 32315913 | 22582 |
| PI 558499 | *A. hypochondriacus* | leaf | 39197764 | 36579 |
| PI 558499 | *A. hypochondriacus* | leaf | 31477634 | 18787 |
| PI 558499 | *A. hypochondriacus* | seedling | 36857940 | 35456 |
| PI 558499 | *A. hypochondriacus* | leaf | 27272930 | 29685 |
| PI 558499 | *A. hypochondriacus* | leaf | 31799201 | 16923 |
| PI 558499 | *A. hypochondriacus* | leaf | 32799196 | 24569 |
| PI 558499 | *A. hypochondriacus* | seedling | 38503380 | 30749 |
| PI 604581 | *A. hypochondriacus* | leaf | 34501001 | 31206 |
| PI 604581 | *A. hypochondriacus* | seedling | 32051291 | 20056 |
| PI 604587 | *A. hypochondriacus* | leaf | 13469060 | 15592 |
| PI 604587 | *A. hypochondriacus* | seedling | 19992232 | 25289 |
| PI 643070 | *A. hypochondriacus* | seedling | 30173461 | 30695 |
| PI 643070 | *A. hypochondriacus* | leaf | 32436428 | 32615 |
| PI 643070 | *A. hypochondriacus* | seedling | 7483949 | 9187 |
| PI 490466 | *A. quitensis* | seedling | 46005628 | 39647 |
| PI 490466 | *A. quitensis* | leaf | 23810641 | 18909 |
| PI 511745 | *A. quitensis* | leaf | 42416284 | 36863 |
| PI 511745 | *A. quitensis* | seedling | 30666794 | 28014 |
| PI 652426 | *A. quitensis* | leaf | 40889587 | 39656 |
| PI 667158 | *A. quitensis* | leaf | 29416173 | 31427 |
| PI 667158 | *A. quitensis* | seedling | 25003043 | 25489 |
| PI 669836 | *A. quitensis* | leaf | 26293386 | 33185 |

**Table S4.**
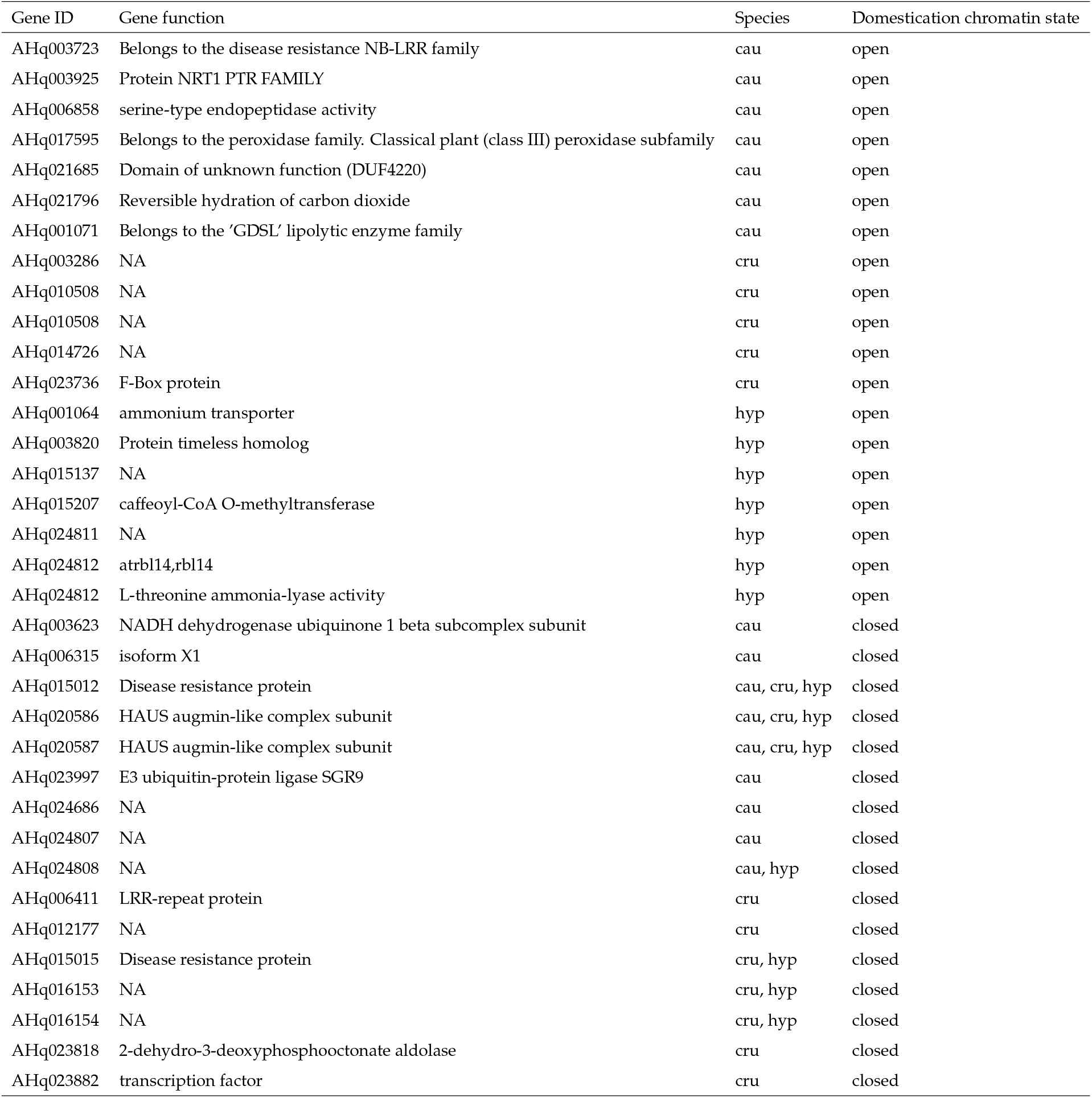
List of genes that were associated with dACRs with selective sweeps.

## References

[1] Joseph R Burger and Trevor S Fristoe. Hunter-gatherer populations inform modern ecology. Proceedings of the National Academy of Sciences, 115(6):1137–1139, 2018.

[2] Jean-Pierre Bocquet-Appel. When the world’s population took off: the springboard of the neolithic demographic transition. Science, 333(6042):560–561, 2011.

[3] Catherine Preece, Alexandra Livarda, Pascal-Antoine Christin, Michael Wallace, Gemma Martin, Michael Charles, Glynis Jones, Mark Rees, and Colin P Osborne. How did the domestication of fertile crescent grain crops increase their yields? Functional ecology, 31(2):387–397, 2017.

[4] Rachel S Meyer and Michael D Purugganan. Evolution of crop species: genetics of domestication and diversification. Nature reviews genetics, 14(12):840–852, 2013.

[5] Anthony Studer, Qiong Zhao, Jeffrey Ross-Ibarra, and John Doebley. Identification of a functional transposon insertion in the maize domestication gene tb1. Nature genetics, 43(11):1160–1163, 2011.

[6] Qiong Lin, Jing Chen, Xuan Liu, Bin Wang, Yaoyao Zhao, Liao Liao, Andrew C Allan, Chongde Sun, Yuquan Duan, Xuan Li, et al. A metabolic perspective of selection for fruit quality related to apple domestication and improvement. Genome Biology, 24(1):95, 2023.

[7] Matthew B Hufford, Xun Xu, Joost Van Heerwaarden, Tanja Pyhäjärvi, Jer-Ming Chia, Reed A Cartwright, Robert J Elshire, Jeffrey C Glaubitz, Kate E Guill, Shawn M Kaeppler, et al. Comparative population genomics of maize domestication and improvement. Nature genetics, 44(7):808–811, 2012.

[8] Gen Xu, Jing Lyu, Qing Li, Han Liu, Dafang Wang, Mei Zhang, Nathan M Springer, Jeffrey Ross-Ibarra, and Jinliang Yang. Evolutionary and functional genomics of dna methylation in maize domestication and improvement. Nature Communications, 11(1):5539, 2020.

[9] Shuai Cao, Kai Chen, Kening Lu, Shiting Chen, Xiyu Zhang, Congcong Shen, Shuangbin Zhu, Yanan Niu, Longjiang Fan, Z Jeffrey Chen, et al. Asymmetric variation in dna methylation during domestication and de-domestication of rice. The Plant Cell, 35(9):3429–3443, 2023.

[10] Alberto Rodriguez-Izquierdo, David Carrasco, Lakshay Anand, Roberta Magnani, Pablo Catarecha, Rosa Arroyo-Garcia, and Carlos M Rodriguez Lopez. Epigenetic differences between wild and cultivated grapevines highlight the contribution of dna methylation during crop domestication. BMC Plant Biology, 24(1):504, 2024.

[11] Yanting Shen, Jixiang Zhang, Yucheng Liu, Shulin Liu, Zhi Liu, Zongbiao Duan, Zheng Wang, Baoge Zhu, Ya-Long Guo, and Zhixi Tian. Dna methylation footprints during soybean domestication and improvement. Genome biology, 19:1–14, 2018.

[12] Shuai Cao, Nunchanoke Sawettalake, Ping Li, Sheng Fan, and Lisha Shen. Dna methylation variations underlie lettuce domestication and divergence. Genome Biology, 25(1):158, 2024.

[13] Paul Bilinski, Patrice S Albert, Jeremy J Berg, James A Birchler, Mark N Grote, Anne Lorant, Juvenal Quezada, Kelly Swarts, Jinliang Yang, and Jeffrey Ross-Ibarra. Parallel altitudinal clines reveal trends in adaptive evolution of genome size in zea mays. PLoS genetics, 14(5):e1007162, 2018.

[14] Lihu Wang, Zhi Luo, Zhiguo Liu, Jin Zhao, Wenping Deng, Hairong Wei, Ping Liu, and Mengjun Liu. Genome size variation within species of chinese jujube (ziziphus jujuba mill.) and its wild ancestor sour jujube (z. acidojujuba cheng et liu). Forests, 10(5):460, 2019.

[15] Concepción M Díez, Brandon S Gaut, Esteban Meca, Enrique Scheinvar, Salvador Montes-Hernandez, Luis E Eguiarte, and Maud I Tenaillon. Genome size variation in wild and cultivated maize along altitudinal gradients. New Phytologist, 199(1):264–276, 2013.

[16] Jonathan F Wendel, Scott A Jackson, Blake C Meyers, and Rod A Wing. Evolution of plant genome architecture. Genome biology, 17:1–14, 2016.

[17] Zaida Vergara and Crisanto Gutierrez. Emerging roles of chromatin in the maintenance of genome organization and function in plants. Genome Biology, 18(1):96, 2017.

[18] Christopher J Hale, Magdalena E Potok, Jennifer Lopez, Truman Do, Ao Liu, Javier Gallego-Bartolome, Scott D Michaels, and Steven E Jacobsen. Identification of multiple proteins coupling transcriptional gene silencing to genome stability in arabidopsis thaliana. PLoS genetics, 12(6):e1006092, 2016.

[19] Agnieszka A Golicz, Prem L Bhalla, David Edwards, and Mohan B Singh. Rice 3d chromatin structure correlates with sequence variation and meiotic recombination rate. Communications biology, 3(1):235, 2020.

[20] Housheng Hansen He, Clifford A Meyer, Hyunjin Shin, Shannon T Bailey, Gang Wei, Qianben Wang, Yong Zhang, Kexin Xu, Min Ni, Mathieu Lupien, et al. Nucleosome dynamics define transcriptional enhancers. Nature genetics, 42(4):343–347, 2010.

[21] Mingkun Huang, Ling Zhang, Limeng Zhou, Wai-Shing Yung, Zhili Wang, Zhixia Xiao, Qianwen Wang, Xin Wang, Man-Wah Li, and Hon-Ming Lam. Identification of the accessible chromatin regions in six tissues in the soybean. Genomics, 114(3):110364, 2022.

[22] Shelby A Blythe and Eric F Wieschaus. Establishment and maintenance of heritable chromatin structure during early drosophila embryogenesis. Elife, 5:e20148, 2016.

[23] Longfei Wang, Guanghong Jia, Xinyu Jiang, Shuai Cao, Z Jeffrey Chen, and Qingxin Song. Altered chromatin architecture and gene expression during polyploidization and domestication of soybean. The Plant Cell, 33(5): 1430–1446, 2021.

[24] Jian Cheng, Xiaoxian Guo, Pengli Cai, Xiaozhi Cheng, Jure Piškur, Yanhe Ma, Huifeng Jiang, and Zhenglong Gu. Parallel evolution of chromatin structure underlying metabolic adaptation. Molecular biology and evolution, 34(11):2870–2878, 2017.

[25] Jingya Yuan, Haojie Sun, Yijin Wang, Lulu Li, Shiting Chen, Wu Jiao, Guanghong Jia, Longfei Wang, Junrong Mao, Zhongfu Ni, et al. Open chromatin interaction maps reveal functional regulatory elements and chromatin architecture variations during wheat evolution. Genome Biology, 23(1):34, 2022.

[26] Cristina M Alexandre, James R Urton, Ken Jean-Baptiste, John Huddleston, Michael W Dorrity, Josh T Cuperus, Alessandra M Sullivan, Felix Bemm, Dino Jolic, Andrej A Arsovski, et al. Complex relationships between chromatin accessibility, sequence divergence, and gene expression in arabidopsis thaliana. Molecular Biology and Evolution, 35 (4):837–854, 2018.

[27] Markus G Stetter, Mireia Vidal-Villarejo, and Karl J Schmid. Parallel seed color adaptation during multiple domestication attempts of an ancient new world grain. Molecular Biology and Evolution, 37(5):1407–1419, 2020.

[28] José Gonçalves-Dias, Akanksha Singh, Corbinian Graf, and Markus G Stetter. Genetic incompatibilities and evolutionary rescue by wild relatives shaped grain amaranth domestication. Molecular Biology and Evolution, 40(8): msad177, 2023.

[29] DJ Lightfoot, David Erwin Jarvis, T Ramaraj, R Lee, EN Jellen, and PJ Maughan. Single-molecule sequencing and Hi-C-based proximity-guided assembly of amaranth (Amaranthus hypochondriacus) chromosomes provide insights into genome evolution. BMC biology, 15:1–15, 2017.

[30] Tom S Winkler, Susanne K Vollmer, Nadine Dyballa-Rukes, Sabine Metzger, and Markus G Stetter. Isoform-resolved genome annotation enables mapping of tissue-specific betalain regulation in amaranth. New Phytologist, 2024.

[31] Stephen I Wright, Newton Agrawal, and Thomas E Bureau. Effects of recombination rate and gene density on transposable element distributions in arabidopsis thaliana. Genome Research, 13(8):1897–1903, 2003.

[32] Damilola A Raiyemo, Jacob S Montgomery, Luan Cutti, Fatemeh Abdollahi, Victor Llaca, Kevin Fengler, Alexander J Lopez, Sarah Morran, Christopher A Saski, David R Nelson, et al. Chromosome-level assemblies of amaranthus palmeri, amaranthus retroflexus, and amaranthus hybridus allow for genomic comparisons and identification of a sex-determining region. The Plant Journal, 121(4):e70027, 2025.

[33] Hengchao Wang, Dong Xu, Sen Wang, Anqi Wang, Lihong Lei, Fan Jiang, Boyuan Yang, Lihua Yuan, Rong Chen, Yan Zhang, et al. Chromosome-scale Amaranthus tricolor genome provides insights into the evolution of the genus Amaranthus and the mechanism of betalain biosynthesis. DNA research, 30(1):dsac050, 2023.

[34] Yong Zhang, Tao Liu, Clifford A Meyer, Jérôme Eeckhoute, David S Johnson, Bradley E Bernstein, Chad Nusbaum, Richard M Myers, Myles Brown, Wei Li, et al. Model-based analysis of ChIP-Seq (MACS). Genome biology, 9(9):1–9, 2008.

[35] Zefu Lu, Alexandre P Marand, William A Ricci, Christina L Ethridge, Xiaoyu Zhang, and Robert J Schmitz. The prevalence, evolution and chromatin signatures of plant regulatory elements. Nature Plants, 5(12):1250–1259, 2019.

[36] Rachel Schwope, Gabriele Magris, Mara Miculan, Eleonora Paparelli, Mirko Celii, Aldo Tocci, Fabio Marroni, Alice Fornasiero, Emanuele De Paoli, and Michele Morgante. Open chromatin in grapevine marks candidate CREs and with other chromatin features correlates with gene expression. The Plant Journal, 107(6):1631–1647, 2021.

[37] Wenbin Mei, Markus G Stetter, Daniel J Gates, Michelle C Stitzer, and Jeffrey Ross-Ibarra. Adaptation in plant genomes. American journal of botany, 105(1):16–19, 2018.

[38] Xianjun Dong, Melissa C Greven, Anshul Kundaje, Sarah Djebali, James B Brown, Chao Cheng, Thomas R Gingeras, Mark Gerstein, Roderic Guigó, Ewan Birney, et al. Modeling gene expression using chromatin features in various cellular contexts. Genome biology, 13:1–17, 2012.

[39] Felix Krueger and Simon R Andrews. Bismark: a flexible aligner and methylation caller for bisulfite-seq applications. bioinformatics, 27(11):1571–1572, 2011.

[40] Shawn J Cokus, Suhua Feng, Xiaoyu Zhang, Zugen Chen, Barry Merriman, Christian D Haudenschild, Sriharsa Pradhan, Stanley F Nelson, Matteo Pellegrini, and Steven E Jacobsen. Shotgun bisulphite sequencing of the arabidopsis genome reveals dna methylation patterning. Nature, 452(7184):215–219, 2008.

[41] Xiaoya Shi, Shuo Cao, Xu Wang, Siyang Huang, Yue Wang, Zhongjie Liu, Wenwen Liu, Xiangpeng Leng, Yanling Peng, Nan Wang, et al. The complete reference genome for grapevine (vitis vinifera l.) genetics and breeding. Horticulture Research, 10(5):uhad061, 2023.

[42] Damon Lisch. Epigenetic regulation of transposable elements in plants. Annual review of plant biology, 60(1):43–66, 2009.

[43] R Keith Slotkin and Robert Martienssen. Transposable elements and the epigenetic regulation of the genome. Nature reviews genetics, 8(4):272–285, 2007.

[44] Sanzhen Liu, Cheng-Ting Yeh, Tieming Ji, Kai Ying, Haiyan Wu, Ho Man Tang, Yan Fu, Dan Nettleton, and Patrick S Schnable. Mu transposon insertion sites and meiotic recombination events co-localize with epigenetic marks for open chromatin across the maize genome. PLoS genetics, 5(11):e1000733, 2009.

[45] Xiao Ma, Fabian E Vaistij, Yi Li, Willem S Jansen van Rensburg, Sarah Harvey, Michael W Bairu, Sonja L Venter, Sydney Mavengahama, Zemin Ning, Ian A Graham, et al. A chromosome-level Amaranthus cruentus genome assembly highlights gene family evolution and biosynthetic gene clusters that may underpin the nutritional value of this traditional crop. The Plant Journal, 107(2):613–628, 2021.

[46] Susanne K Vollmer, Markus G Stetter, and Goetz Hensel. First successful targeted mutagenesis using crispr/cas9 in stably transformed grain amaranth tissue. bioRxiv, pages 2025–02, 2025.

[47] Michelle C Stitzer, Sarah N Anderson, Nathan M Springer, and Jeffrey Ross-Ibarra. The genomic ecosystem of transposable elements in maize. PLoS genetics, 17(10):e1009768, 2021.

[48] Bo Zhu, Wenli Zhang, Tao Zhang, Bao Liu, and Jiming Jiang. Genome-wide prediction and validation of intergenic enhancers in arabidopsis using open chromatin signatures. The Plant Cell, 27(9):2415–2426, 2015.

[49] J Engelhorn et al. Genetic variation at transcription factor binding sites largely explains phenotypic heritability in maize. bioRxiv, pages 2023–2008, 2024.

[50] Zefu Lu, William A Ricci, Robert J Schmitz, and Xiaoyu Zhang. Identification of cis-regulatory elements by chromatin structure. Current opinion in plant biology, 42:90–94, 2018.

[51] Eli Rodgers-Melnick, Daniel L Vera, Hank W Bass, and Edward S Buckler. Open chromatin reveals the functional maize genome. Proceedings of the National Academy of Sciences, 113(22):E3177–E3184, 2016.

[52] Yuting Liu, Xiang Gao, Hongjun Liu, Xuerong Yang, Xiao Liu, Fang Xu, Yuzhi Zhu, Qingyun Li, Liangliang Huang, Fang Yang, et al. Constraint of accessible chromatins maps regulatory loci involved in maize speciation and domestication. Nature Communications, 16(1):2477, 2025.

[53] Zachary H Lemmon, Robert Bukowski, Qi Sun, and John F Doebley. he role of cis regulatory evolution in maize domestication. PLoS genetics, 10(11):e1004745, 2014.

[54] Ryan A Rapp, Candace H Haigler, Lex Flagel, Ran H Hovav, Joshua A Udall, and Jonathan F Wendel. Gene expression in developing fibres of upland cotton (gossypium hirsutum l.) was massively altered by domestication. BMC biology, 8:1–15, 2010.

[55] Christopher Sauvage, Andrea Rau, Charlotte Aichholz, Joel Chadoeuf, Gautier Sarah, Manuel Ruiz, Sylvain Santoni, Mathilde Causse, Jacques David, and Sylvain Glémin. Domestication rewired gene expression and nucleotide diversity patterns in tomato. The Plant Journal, 91(4):631–645, 2017.

[56] C Darwin. The variation of plants and animals under domestication. revised 1875. Murray: London, 1868.

[57] Ian K Greaves, Michael Groszmann, Aihua Wang, W James Peacock, and Elizabeth S Dennis. Inheritance of trans chromosomal methylation patterns from arabidopsis f1 hybrids. Proceedings of the National Academy of Sciences, 111 (5):2017–2022, 2014.

[58] Stéphanie Durand, Nicolas Bouché, Elsa Perez Strand, Olivier Loudet, and Christine Camilleri. Rapid establishment of genetic incompatibility through natural epigenetic variation. Current Biology, 22(4):326–331, 2012.

[59] Juntao Hu, Sara J Smith, Tegan N Barry, Heather A Jamniczky, Sean M Rogers, and Rowan DH Barrett. Heritability of dna methylation in threespine stickleback (gasterosteus aculeatus). Genetics, 217(1):iyab001, 2021.

[60] Brieanne Vaillancourt and C Robin Buell. High molecular weight dna isolation method from diverse plant species for use with oxford nanopore sequencing. BioRxiv, page 783159, 2019.

[61] Wouter De Coster and Rosa Rademakers. Nanopack2: population-scale evaluation of long-read sequencing data. Bioinformatics, 39(5):btad311, 2023.

[62] Haoyu Cheng, Gregory T Concepcion, Xiaowen Feng, Haowen Zhang, and Heng Li. Haplotype-resolved de novo assembly using phased assembly graphs with hifiasm. Nature methods, 18(2):170–175, 2021.

[63] Heng Li and Richard Durbin. ast and accurate long-read alignment with burrows–wheeler transform. Bioinformatics, 26(5):589–595, 2010.

[64] Chenxi Zhou, Shane A McCarthy, and Richard Durbin. Yahs: yet another hi-c scaffolding tool. Bioinformatics, 39(1): btac808, 2023.

[65] Neva C Durand, James T Robinson, Muhammad S Shamim, Ido Machol, Jill P Mesirov, Eric S Lander, and Erez Lieberman Aiden. Juicebox provides a visualization system for hi-c contact maps with unlimited zoom. Cell systems, 3(1):99–101, 2016.

[66] Yu Chen, Yixin Zhang, Amy Y Wang, Min Gao, and Zechen Chong. Accurate long-read de novo assembly evaluation with inspector. Genome Biology, 22:1–21, 2021.

[67] Jullien M Flynn, Robert Hubley, Clément Goubert, Jeb Rosen, Andrew G Clark, Cédric Feschotte, and Arian F Smit. Repeatmodeler2 for automated genomic discovery of transposable element families. Proceedings of the National Academy of Sciences, 117(17):9451–9457, 2020.

[68] Nansheng Chen. Using repeat masker to identify repetitive elements in genomic sequences. Current protocols in bioinformatics, 5(1):4–10, 2004.

[69] Lars Gabriel, Tom Brů na, Katharina J Hoff, Matthis Ebel, Alexandre Lomsadze, Mark Borodovsky, and Mario Stanke. BRAKER3: Fully automated genome annotation using RNA-Seq and protein evidence with GeneMark-ETP, AUGUSTUS and TSEBRA. Biorxiv, 2023.

[70] Evgenia V Kriventseva, Dmitry Kuznetsov, Fredrik Tegenfeldt, Mosè Manni Renata Dias, Felipe A Simão, and Evgeny M Zdobnov. OrthoDB v10: sampling the diversity of animal, plant, fungal, protist, bacterial and viral genomes for evolutionary and functional annotations of orthologs. Nucleic Acids Research, 47(D1):D807–D811, 2019.

[71] JW Clouse, D Adhikary, JT Page, T Ramaraj, MK Deyholos, JA Udall, DJ Fairbanks, EN Jellen, and PJ Maughan. The amaranth genome: genome, transcriptome, and physical map assembly. The Plant Genome, 9(1):plantgenome2015– 07, 2016.

[72] Alexander Dobin and Thomas R Gingeras. Mapping RNA-seq reads with STAR. Current Protocols in Bioinformatics, 51(1):11–14, 2015.

[73] Heng Li. Minimap2: pairwise alignment for nucleotide sequences. Bioinformatics, 34(18):3094–3100, 2018.

[74] Manuel Tardaguila, Lorena de la Fuente, Cristina Marti, Cécile Pereira, Francisco Jose Pardo-Palacios, Hector del Risco, Marc Ferrell, Maravillas Mellado, Marissa Macchietto, Kenneth Verheggen, et al. SQANTI: extensive characterization of long-read transcript sequences for quality control in full-length transcriptome identification and quantification. Genome Research, 28(3):396–411, 2018.

[75] Shiyuyun Tang, Alexandre Lomsadze, and Mark Borodovsky. Identification of protein coding regions in RNA transcripts. Nucleic acids research, 43(12):e78–e78, 2015.

[76] Lars Gabriel, Katharina J Hoff, Tom Brů na, Mark Borodovsky, and Mario Stanke. TSEBRA: Transcript Selector for BRAKER. BMC bioinformatics, 22(1):1–12, 2021.

[77] Felipe A Simão, Robert M Waterhouse, Panagiotis Ioannidis, Evgenia V Kriventseva, and Evgeny M Zdobnov. BUSCO: assessing genome assembly and annotation completeness with single-copy orthologs. Bioinformatics, 31 (19):3210–3212, 2015.

[78] Rainer Schwacke, Gabriel Y Ponce-Soto, Kirsten Krause, Anthony M Bolger, Borjana Arsova, Asis Hallab, Kristina Gruden, Mark Stitt, Marie E Bolger, and Björn Usadel. Mapman4: a Refined Protein Classification and Annotation Framework Applicable to Multi-Omics Data Analysis. Molecular plant, 12(6):879–892, 2019.

[79] Carlos P Cantalapiedra, Ana Hernández-Plaza, Ivica Letunic, Peer Bork, and Jaime Huerta-Cepas. eggNOGmapper v2: Functional Annotation, Orthology Assignments, and Domain Prediction at the Metagenomic Scale. Molecular Biology and Evolution, 38(12):5825–5829, 2021.

[80] Shujun Ou, Weija Su, Yi Liao, Kapeel Chougule, Jireh RA Agda, Adam J Hellinga, Carlos Santiago Blanco Lugo, Tyler A Elliott, Doreen Ware, Thomas Peterson, et al. Benchmarking transposable element annotation methods for creation of a streamlined, comprehensive pipeline. Genome biology, 20:1–18, 2019.

[81] David M Emms and Steven Kelly. Orthofinder: phylogenetic orthology inference for comparative genomics. Genome biology, 20:1–14, 2019.

[82] Damilola A Raiyemo, Luan Cutti, Eric L Patterson, Victor Llaca, Kevin Fengler, Jacob S Montgomery, Sarah Morran, Todd A Gaines, and Patrick J Tranel. A phased chromosome-level genome assembly provides insights into the evolution of sex chromosomes in amaranthus tuberculatus. bioRxiv, pages 2024–05, 2024.

[83] J Mitchell McGrath, Andrew Funk, Paul Galewski, Shujun Ou, Belinda Townsend, Karen Davenport, Hajnalka Daligault, Shannon Johnson, Joyce Lee, Alex Hastie, et al. A contiguous de novo genome assembly of sugar beet el10 (beta vulgaris l.). DNA research, 30(1):dsac033, 2023.

[84] Elodie Rey, Peter J Maughan, Florian Maumus, Daniel Lewis, Leanne Wilson, Juliana Fuller, Sandra M Schmöckel, Eric N Jellen, Mark Tester, and David E Jarvis. A chromosome-scale assembly of the quinoa genome provides insights into the structure and dynamics of its subgenomes. Communications Biology, 6(1):1263, 2023.

[85] Jacob Montgomery, Sarah Morran, Dana R MacGregor, J Scott McElroy, Paul Neve, Célia Neto, Martin M Vila-Aiub, Maria Victoria Sandoval, Analia I Menéndez, Julia M Kreiner, et al. Current status of community resources and priorities for weed genomics research. Genome biology, 25(1):139, 2024.

[86] Hengchi Chen, Arthur Zwaenepoel, and Yves Van de Peer. wgd v2: a suite of tools to uncover and date ancient polyploidy and whole-genome duplication. Bioinformatics, 40(5):btae272, 2024.

[87] Yupeng Wang, Haibao Tang, Jeremy D DeBarry, Xu Tan, Jingping Li, Xiyin Wang, Tae-ho Lee, Huizhe Jin, Barry Marler, Hui Guo, et al. Mcscanx: a toolkit for detection and evolutionary analysis of gene synteny and collinearity. Nucleic acids research, 40(7):e49–e49, 2012.

[88] Joselyn J Todd and Lila O Vodkin. Duplications that suppress and deletions that restore expression from a chalcone synthase multigene family. The Plant Cell, 8(4):687–699, 1996.

[89] Fu-Xiang Wang, Guan-Dong Shang, Lian-Yu Wu, Yan-Xia Mai, Jian Gao, Zhou-Geng Xu, and Jia-Wei Wang. Protocol for assaying chromatin accessibility using ATAC-seq in plants. STAR protocols, 2(1):100289, 2021.

[90] Md Vasimuddin, Sanchit Misra, Heng Li, and Srinivas Aluru. Efficient architecture-aware acceleration of BWA-MEM for multicore systems. In 2019 IEEE international parallel and distributed processing symposium (IPDPS), pages 314–324. IEEE, 2019.

[91] Broad Institue. Picard tools—by broad institute, 2018.

[92] Zuguang Gu, Lei Gu, Roland Eils, Matthias Schlesner, and Benedikt Brors. circlize implements and enhances circular visualization in R. Bioinformatics, 30:2811–2812, 2014.

[93] Lihua J Zhu, Claude Gazin, Nathan D Lawson, Hervé Pagès, Simon M Lin, David S Lapointe, and Michael R Green. ChIPpeakAnno: a Bioconductor package to annotate ChIP-seq and ChIP-chip data. BMC bioinformatics, 11:1–10, 2010.

[94] Matthew D Young, Matthew J Wakefield, Gordon K Smyth, and Alicia Oshlack. Gene ontology analysis for RNA-seq: accounting for selection bias. Genome Biology, 11:R14, 2010.

[95] Nicolas L Bray, Harold Pimentel, Páll Melsted, and Lior Pachter. Near-optimal probabilistic rna-seq quantification. Nature biotechnology, 34(5):525–527, 2016.

[96] R Core Team. R: A Language and Environment for Statistical Computing. R Foundation for Statistical Computing, Vienna, Austria, 2023. URL https://www.R-project.org/.

[97] Michael Lawrence, Wolfgang Huber, Hervé Pagès, Patrick Aboyoun, Marc Carlson, Robert Gentleman, Martin Morgan, and Vincent Carey. Software for computing and annotating genomic ranges. PLoS Computational Biology, 9, 2013. doi: 10.1371/journal.pcbi.1003118. URL http://www.ploscompbiol.org/article/info%3Adoi%2F10.1371%2Fjournal.pcbi.1003118.

[98] R Core Team. R: A Language and Environment for Statistical Computing. R Foundation for Statistical Computing, Vienna, Austria, 2024. URL https://www.R-project.org/.

[99] José Gonçalves-Dias and Markus G Stetter. Popamaranth: a population genetic genome browser for grain amaranths and their wild relatives. G3, 11(7):jkab103, 2021.

[100] Nikolaos Alachiotis and Pavlos Pavlidis. Raisd detects positive selection based on multiple signatures of a selective sweep and snp vectors. Communications biology, 1(1):79, 2018.

[101] Aaron R Quinlan and Ira M Hall. Bedtools: a flexible suite of utilities for comparing genomic features. Bioinformatics, 26(6):841–842, 2010.

